# Endoparasitoid lifestyle promotes endogenization and domestication of dsDNA viruses

**DOI:** 10.1101/2022.11.16.515002

**Authors:** Benjamin Guinet, David Lepetit, Sylvain Charlat, Peter N Buhl, David G Notton, Astrid Cruaud, Jean-Yves Rasplus, Julia Stigenberg, Damien M. de Vienne, Boussau Bastien, Julien Varaldi

**Affiliations:** Université Lyon 1, CNRS, Laboratoire de Biométrie et Biologie Evolutive UMR 5558, F-69622 Villeurbanne, France; Zoological Museum, Department of Entomology, University of Copenhagen, Universitetsparken 15, DK-2100 Copenhagen, Denmark; Natural Sciences Department, National Museums Collection Centre, 242 West Granton Road, Granton, Edinburgh, EH5 1JA, United Kingdom; Department of Zoology, Swedish Museum of Natural History, Box 50007, 104 05 Stockholm, Sweden; INRAE, UMR 1062 CBGP, 755 avenue du campus Agropolis CS 30016, 34988 Montferrier-sur-Lez, France

**Author notes:** Authors correspondence (BG); (JV).

## Abstract

The accidental endogenization of viral elements within eukaryotic genomes can occasionally provide significant evolutionary benefits, giving rise to their long-term retention, that is, to viral domestication. For instance, in some endoparasitoid wasps (whose immature stages develop inside their hosts), the membrane-fusion property of double-stranded DNA viruses has been repeatedly domesticated following ancestral endogenizations. The endogenized genes provide female wasps with a delivery tool to inject virulence factors that are essential to the developmental success of their offspring. Because all known cases of viral domestication involve endoparasitic wasps, we hypothesized that this lifestyle, relying on a close interaction between individuals, may have promoted the endogenization and domestication of viruses. By analyzing the composition of 124 Hymenoptera genomes, spread over the diversity of this clade and including free-living, ecto- and endoparasitoid species, we tested this hypothesis. Our analysis first revealed that double-stranded DNA viruses, in comparisons with other viral genomic structures (ssDNA, dsRNA, ssRNA), are more often endogenized and domesticated (that is, retained by selection) than expected from their estimated abundance in insect viral communities. Secondly, our analysis indicates that the rate at which dsDNA viruses are endogenized is higher in endoparasitoids than in ectoparasitoids or free-living hymenopterans, which also translates into more frequent events of domestication. Hence, these results are consistent with the hypothesis that the endoparasitoid lifestyle has facilitated the endogenization of dsDNA viruses, in turn increasing the opportunities of domestications that now play a central role in the biology of many endoparasitoid lineages.

## Introduction

The recent boom of genome sequencing programs has revealed the abundance of DNA fragments of viral origin within eukaryotic genomes. These so-called Endogenous Viral Elements (EVEs) stem from endogenization events that not only involve retroviruses as donors (as could be expected from their natural lifecycle) but also viruses that do not typically integrate into their host chromosomes [1, 2, 3]. In insects, where retroviruses have yet to be found, endogenization events have involved various non-retroviral viruses: three families of large double-stranded (ds) DNA viruses, at least 22 families of RNA viruses, and three families of single-stranded (ss) DNA viruses [4]. Degeneracy and loss is likely the fate of most EVEs, since they do not *a priori* benefit their hosts. Still, several studies have reported that EVEs can be retained by selection, thus becoming *domesticated* [5]. The functions involved include defensive properties against related viruses in mosquitoes [6, 7], against macroparasites in some Lepidoptera [8], or modifications in the expression of genes involved in dispersal in aphids [9]. Beyond insects, the membrane fusion capacity of viruses, allowing their entry into host cells, has been repeatedly co-opted in three metazoan clades: mammals, viviparous lizards and parasitoid wasps. In placental mammals and viviparous Scincidae lizards, domestication of the *syncytin* protein from retroviruses has allowed the emergence of the placenta, through the development of the syncytium (composed of fused cells) involved in metabolic exchanges between the mother and the fetus [10, 11]. A similar fusogenic property was repeatedly co-opted by parasitoids belonging to the Hymenoptera order through the endogenization and domestication of complex viral machineries deriving from large dsDNA viruses [12, 4]. The numerous retained viral genes allow parasitoid wasps to produce virus-like structures (VLS) within their reproductive apparatus. These are injected into the wasp’s host, together with their eggs, and protect the wasp progeny against the host immune response. This protection is achieved thanks to the ability of VLS to deliver virulence factors in the form of genes (in which case VLS are called polydnavirus - PDV) or proteins (in which case VLS are called Virus-like particles - VLPs) to host immune cells (reviewed in [13, 14]). So far, 5 independent cases of such viral domestication have been detected in parasitoid wasps, four of them falling within the Ichneumonoidea superfamily [15, 16, 17, 18] and one in the Cynipoidea superfamily [19]. The four cases where the donor virus family has been unequivocally identified point towards dsDNA viruses. More specifically, the domesticated EVEs (hereafter, dEVEs) derive from the *Nudiviridae* family in three cases [15, 17, 18] while the forth involves a putatively new viral family denoted “LbFV-like” [19]. Notably, all these domestication events took place in endoparasitoids, that is, in species that deposit their eggs inside the hosts, as opposed to ectoparasitoids that lay on their surface.

Beyond these well characterized events of viral domestication in Hymenoptera, additional cases of endogenization have been uncovered, in studies that enlarged the taxonomic focus of either the hosts [20, 21, 22] or the viruses that were considered [23, 20, 24, 25]. Here, we complement this earlier work by expanding the range of both the hosts and viruses under study, and by further analyzing which endogenization cases have been followed by a domestication event.

To this end, we developed a bioinformatic pipeline to detect endogenization events involving any kind of viruses (DNA/RNA, single-stranded, double-stranded), at the scale of the whole Hymenoptera order. This analysis first allowed us to test whether the propensity of viruses to enter Hymenoptera genomes, and to be domesticated, depend on their genomic structure (in line with the pattern observed so far, where only dsDNA viruses have been involved in domestication events as described above). We then tested whether the lifestyle of the species (free-living, endoparasitoid, ectoparasitoid) correlates with their propensity to integrate and domesticate viruses. Our working hypothesis was that the endoparasitoid lifestyle may be associated with a higher rate of viral endogenization and / or a higher rate of domestication events, for two non-exclusive reasons related either to the exposure to new viruses and the adaptive value of the endogenized elements.

First, a higher endogenization rate may simply stem from a higher exposure to viruses. Such an effect could be at play in endoparasitoids due to the intimate interaction between the parasitoid egg or larva and the host. In other words, the endo-parasitic way of life may facilitate the acquisition of new viruses deriving from the hosts. Notably, this lifestyle may also facilitate the maintenance and spread of newly acquired viruses within wasp populations. Indeed, endoparasitoid wasps often inject not only eggs but also venomic compounds (typically produced in the venom gland or in calyx cells) where viruses can be present and may thus be vertically transmitted [26]). In addition, the confinement of the several developing wasps within a single host may facilitate viral horizontal transmission and its subsequent spread in wasps populations (e.g. [27]).

Second, a higher rate of domestication in endoparasitoids may be the consequence of a particular selective regime. This is expected since, these insects are facing the very special challenge of resisting the host immune system, contrary to other lifestyles. This selective pressure may promote the co-option of viral functions such as the above-mentioned membrane fusion activity, that provide a very effective mean to deliver virulence factors.

Our analysis reveals numerous new instances of endogenization events, some of which are also characterized by signatures of molecular domestication. We found a clear enrichment in endogenization events deriving from dsDNA viruses as compared to those with other genomic structures. While the data did not reveal a significant effect of Hymenoptera lifestyles on the acquisition of dsRNA, ssRNA or ssDNA viruses, it supports the hypothesis that genes from dsDNA viruses are more often endogenized and domesticated in endoparasitoids than in free-living and ectoparasitoid species.

## Results

We screened for EVEs 124 Hymenoptera genome assemblies, including 24 ectoparasitoids, 37 endoparasitoids and 63 free-living species (the list can be found on the github repository under the name : Assembly_genome_informations.csv). EVEs were identified using a sequence-homology approach based on a comprehensive viral protein database. Different confidence levels (ranging from A to D) were associated with the various EVEs inferred, where the A score indicates a maximal confidence level for endogenization. This confidence index is based upon sequencing depth combined with the presence of eukaryotic genes and/or transposable elements in the genomic environment of the candidate loci (as detailed in the MM section). By default, the four categories are included in the analysis, but unless otherwise stated statistical tests based on the A category only led to the same conclusions (see FigureS13-7 for more details). Our analysis further included an inference of the phylogenetic relationships among homologous EVEs, that was used to map endogenization events on the Hymenoptera species tree. Finally, inferences of domestication events relied upon signatures of purifying selection in the integrated genes (based on dN/dS estimates) and/or on expression data.

An important objective of our analysis is to detect and enumerate not only endogenous viral elements (EVEs) but also endogenization *events* that can explain the presence of these EVEs. Indeed, an EVE denotes a single gene of viral origin in a single species. Several neighboring EVEs in a genome most likely result from the endogenization of a single viral genome, and homologous EVEs shared by several closely related species may further stem from a single ancestral endogenization events. This distinction is critical when it comes to examining the effect of various factors on the probability of integrating EVEs, which implies counting events rather than EVEs. As an example, consider the *Leptopilina* case [19], involving 13 EVEs shared by 3 closely related species. In this wasp genus, based on previous findings, we expect the 39 EVEs to be grouped into a single endogenization event. Our pipeline appropriately detected 36 EVEs (out of 39) and correctly aggregated them into a single endogenization event mapped on the branch leading to the *Leptopilina* genus. Furthermore, because some of the genes involved are inferred as domesticated, this event is appropriately classified as a domestication event (see Figure 1 and Figure S14 for more canonical examples). In total, the pipeline correctly detected 88.4% (152/172) of the EVEs involved in our four “positive controls”, previously described as mediating the protection of young wasps against their host immune system. Among them, 71.82% were inferred as being domesticated. Out of the 152 positive controls EVEs, 147 were grouped into 4 independent endogenization events, as was expected. The remaining 5 genes had peculiar histories that led our pipeline to infer two additional spurious events (Table S1). All detailed results regarding EVEs and dEVEs can be found on the GitHub repository under the name : All_EVEs_dEVEs_informations.txt.

**Figure 1.**
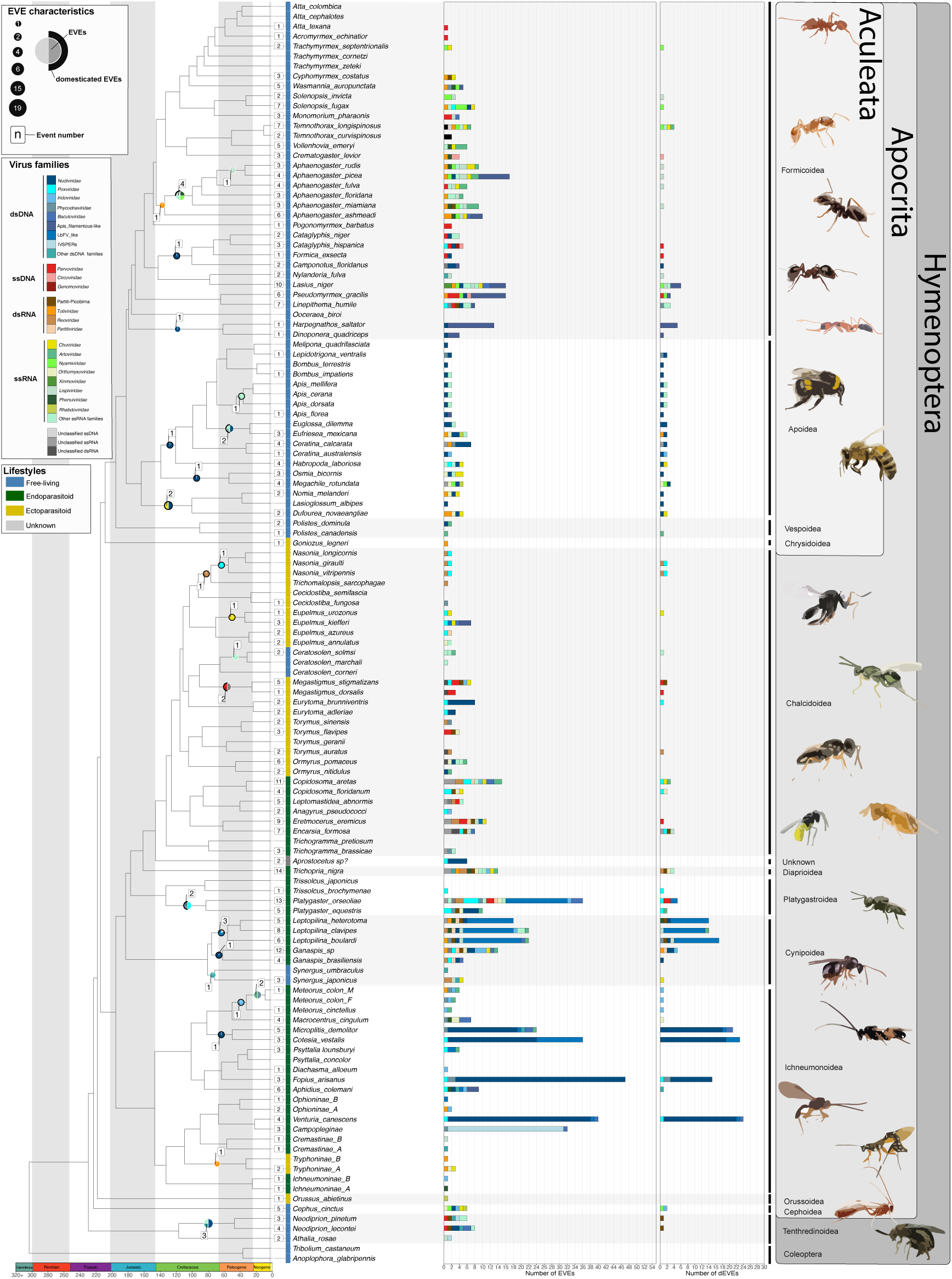
Endogenous Viral Elements and their domestication status in Hymenoptera. Lifestyles are displayed next to species names (blue: free-living, green: endoparasitoid, yellow: ectoparasitoid, grey: unknown). The number of EVEs and domesticated EVEs (dEVEs) found in each species are represented respectively by the first and second facet of the horizontal histograms. Colors along these histograms indicate the potential donor viral families (where blue tones correspond to viral dsDNA viruses, red tones to ssDNA viruses, orange/brown tones to dsRNA viruses and green tones to ssRNA genomes). Endogenous Viral elements (EVEs) shared by multiple species and classified within the same event are represented by circles whose size is proportional to their number; those that are considered as domesticated (dEVEs) are surrounded by a black border. Numbers in the white boxes correspond to the number of endogenization events inferred. As an example, *Megastigmus dorsalis* and *Megastigmus stigmatizans* are ectoparasitoids (yellow) sharing a common endogenization event (within the Cluster21304) that likely originated from an unclassified dsRNA virus (grey color in circle), and shows no sign of domestication (no black border around the grey part of the circle). The figure was inspired from the work of [28]. Details on the phylogenetic inference and time calibration can be found in the MM section; bootstrap information can be found in TableS2; details on lifestyle assignation can be found on the github repository under the name : Assembly_genome_informations.csv

**Figure 2.**
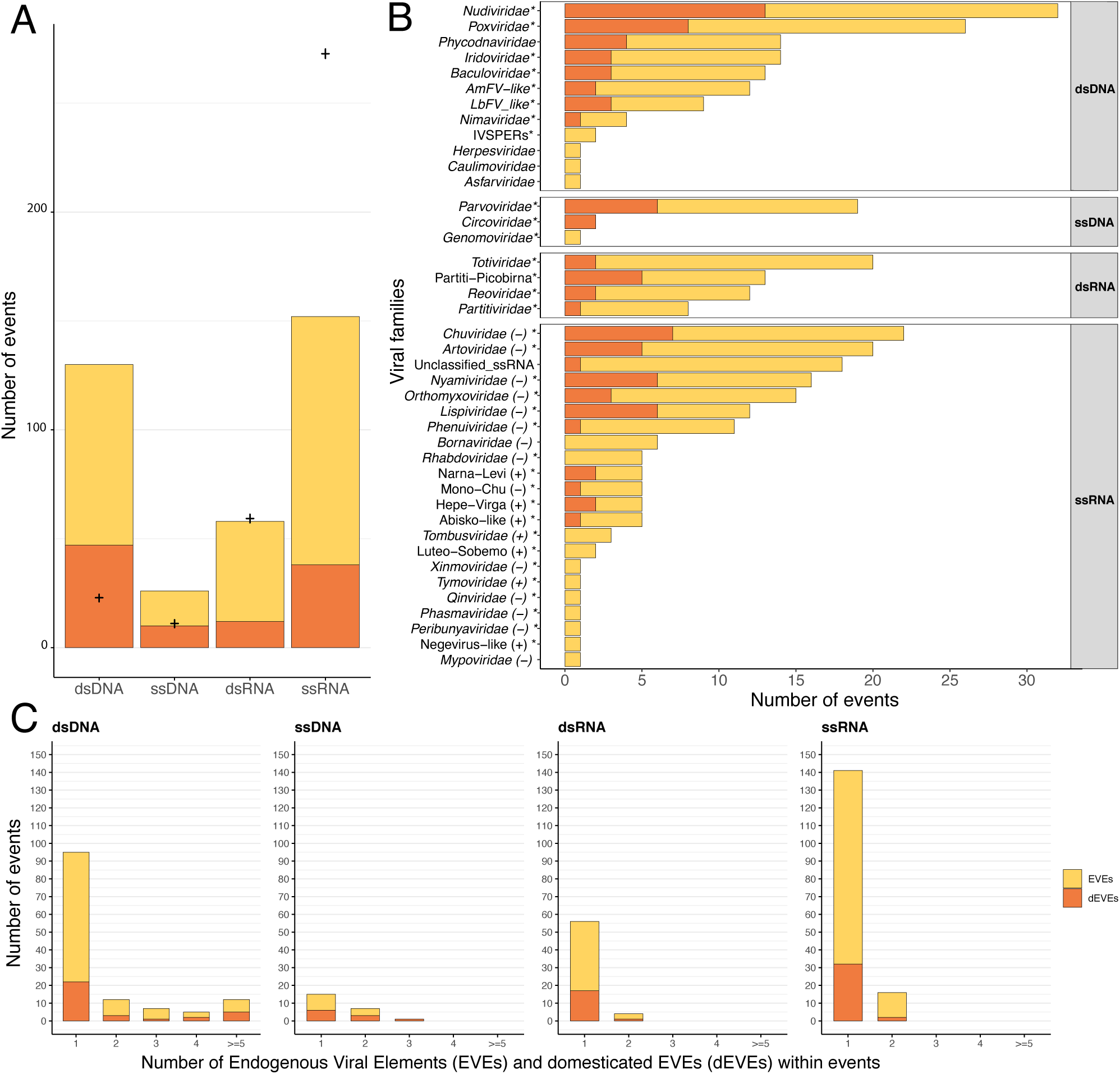
Endogenization involves all types of viral genomic structures. In all three panels, a yellow color indicates endogenization events that have not been followed by domestication, while orange indicates domestication events. **A** : Distribution of the number of events inferred, according to the four categories of viral genomic structures. The crosses refer to the expected number of endogenization events for each category based on its estimated relative abundance in insects (see details in Materials and methods and virus-infecting data in Excel tab:All_virus_infecting_insects_informations). **B** : Distribution of the various viral families involved in endogenization events. The polarity of ssRNA viruses is displayed next to the family name. Events involving multiple putative families (i.e. where several viral families are present in the same scaffold) have been excluded from the count. The star next to the family name indicates that the viral family is known to infect insects. **C** : Distribution of the number of EVEs per event across viral categories.

### Endogenizations involve all viral genomic structures

A total of 1261 EVEs have been inferred in the whole dataset (TableS2,Figure1). These come from 367 endogenization events, the majority of which involved ssRNA and dsDNA viruses (41% and 35%, respectively) (TableS2). Among the tip branches leading to the 124 species under study, 113 underwent one endogenization event, with a maximum of 14 events and a median of 3 (Figure1). In total, 91% of the events (331) occurred on tip branches, and the remaining 9% are shared by at least two closely related species (TableS2, Figure1). To assess the validity of the procedure used to aggregate multiple EVEs into a single shared ancestral endogenization, we assessed whether EVEs inferred as homologous shared a common genomic environment. We thus tested for the presence of homologous loci in descendant species around the shared ancestral EVEs (using blastn searches between the corresponding scaffolds, see details in Materials and methods). Among the 36 endogenization events that involved at least two species, 31 were found to carry more homologous loci around the insertion sites than expected by chance (see details in Materials and methods). Notably, the majority of endogenization events involved a single EVE (a single gene) and only 12 (all from dsDNA viruses), involved the concomitant integration of more than 4 viral genes (Figure2-C).

A total of 40 different viral clades (usually families) were inferred as putative donors. Most of them (34) are known to infect insects (Figure2-B) and these account for the majority of the 331 endogenization events. However, we found 36 EVEs (24 endogenization events), including 20 high-confidence ones (A-ranked), that derived from 6 viral families not previously reported to infect insects (*Phycodnaviridae*, *Herpesviridae*, *Caulimoviridae*, *Asfaviridae*, *Bornaviridae* and *Mypoviridae*). However, in those cases, the true viral donors may belong to unknown clades that do infect insects. Indeed, although the homology with viral proteins was convincing (median e-value was 9.4095e-12 [min = 9.212e-129, max = 3.305e-08]), the average percentage identity was relatively low (38% [min = 23.2%, max = 79.1%], suggesting that these loci may originate from unknown viruses that are only distantly related to their closest relatives in public databases.

### Double-stranded DNA viruses are over-represented in endogenization events

Most of the endogenization events recorded involve ssRNA and dsDNA viruses. But do these proportions simply mirror the diversity and respective abundances of the different kinds of viruses encountered by insects? The analysis summarized in (Figure2-A) (see details in Materials and methods) indicates this is not the case. More specifically, it shows that dsDNA viruses are more frequently endogenized than expected on the basis of their representation in the databases, while ssRNA viruses are under-represented (*x*^2^ = 213.36 and 221.38, respectively, for endogenization events and domestication events, d.f. = 3, both p-value < 2.2e-16). Notably, this result is not purely driven by the presence in our data set of the four positive controls (previously described cases of viral domestication, that all involve dsDNA viruses as donors). Finally, among endogenization events involving ssRNA viruses, we found an over-representation of negative stranded ssRNA compared to their relative abundance in public databases (72.2% compared to 32.6% in the databases, *x*^2^=145.87, d.f.=1, p-value < 2.2e-16; see supplemental information for a discussion).

### Endogenizations of dsDNA viruses are more frequent in endoparasitoid species

Next, we sought to characterize the factors that could explain the patterns of endogenization events inferred (Figure1). To this end, for each viral genomic structure, we assessed whether endogenization events were evenly distributed among the three wasp lifestyles, taking into account their respective frequencies in the dataset. No significant departure from the null hypothesis was detected for endogenization involving ssDNA, dsRNA or ssRNA viruses (Fisher exact test p-values BH corrected > 0.05). On the contrary, we detected a highly significant enrichment of dsDNA viruses endogenization events in endoparasitoid species, and conversely a deficiency in free-living and ectoparasitoid species (corrected p-value = 7.8e-04, Figure 3-A).

**Figure 3.**
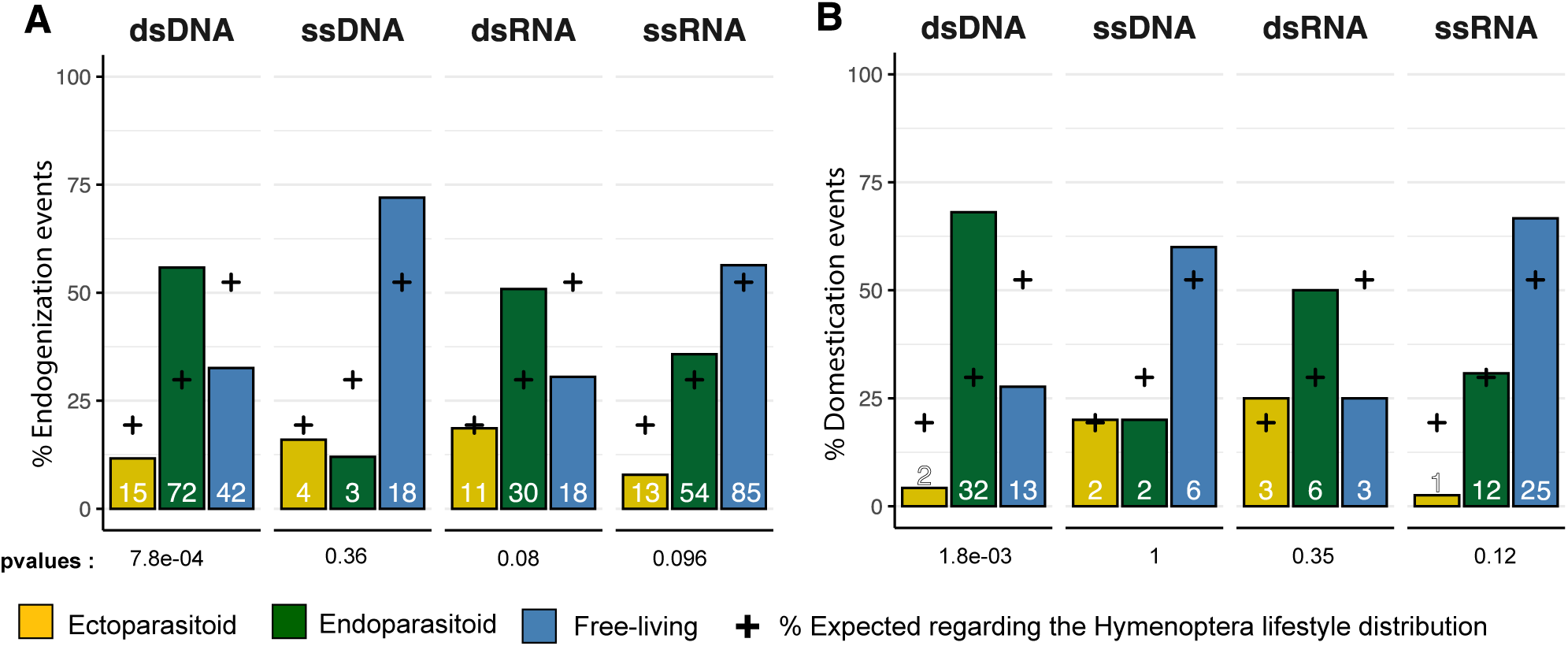
Endogenization and domestication of dsDNA viruses are most prevalent in endoparasitoid species. **A**: Distribution of viral endogenization events (Event) and **B** of domestication events (dEVEs) across Hymenoptera lifestyles. Crosses indicate the expected proportion of events associated with the different lifestyles, based on the respective frequencies in our database (ectoparasitoid = 24/124, endoparasitoid = 37/124, free-living = 63/124). The p-values are the results of Fisher’s tests comparing the observed and expected distributions. Numbers inside the bars indicate the absolute numbers of events inferred. The ancestral states of the nodes, in terms of lifestyle, were inferred in a Bayesian analysis (see details in Materials and methods).

**Figure 4.**
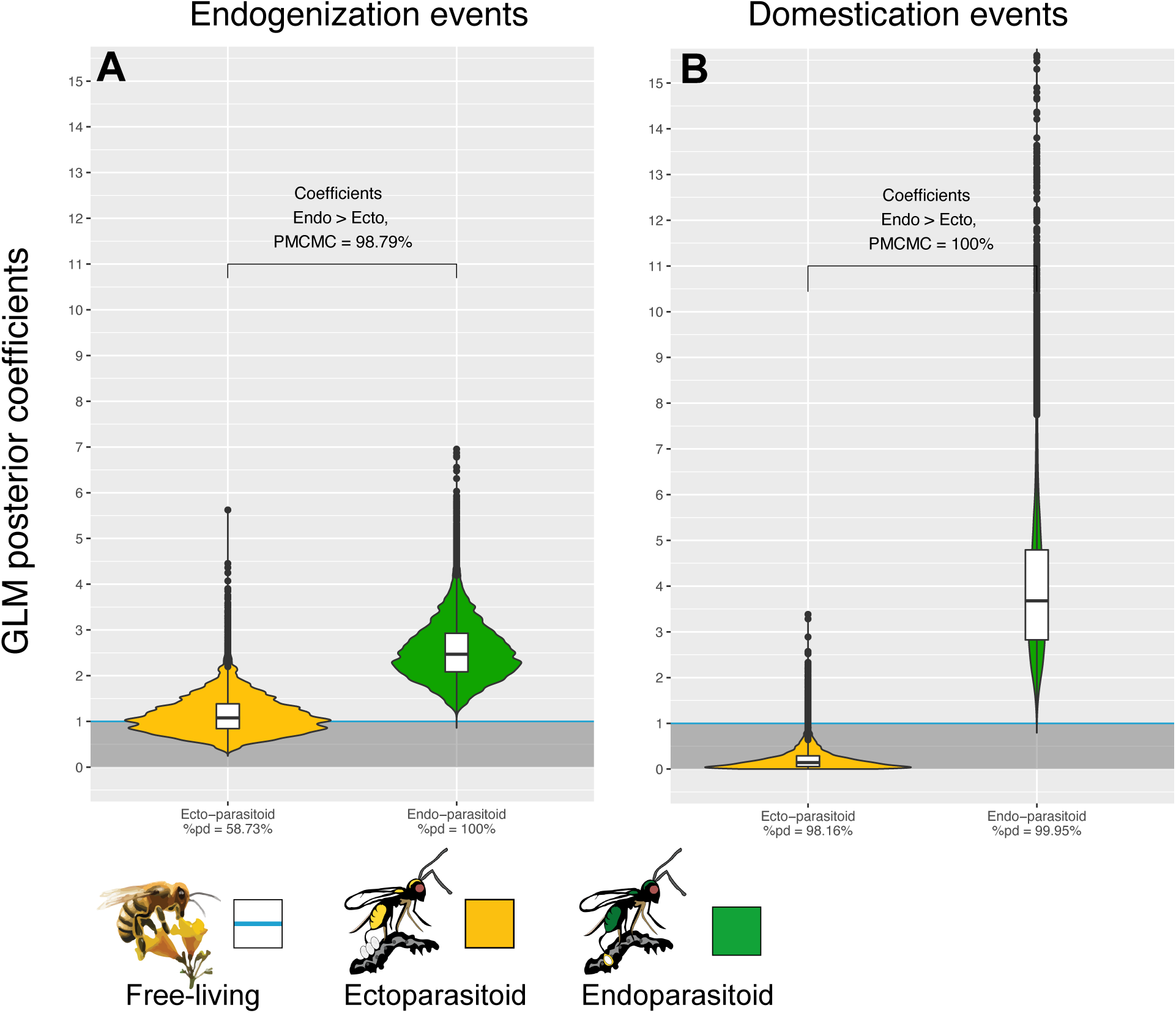
Endogenization and domestication of dsDNA viruses are more frequent in endoparasitoid species. Violin plots represent the posterior distribution of the coefficients obtained under the different GLM models (after exponential transformation to obtain a rate relative to free-living species). The coefficients are derived from 1000 independent GLM models, where 1000 probable scenarios of ancestral states at nodes were sampled randomly among the MCMCM iterations (see details in Materials and methods). Branches from nodes older than 160 million years were removed from the dataset. The %pd is the probability of direction and indicates the proportion of the posterior distribution where the coefficients have the same sign as the median coefficient. *P_MCMCM_* indicates the proportion of MCMC iterations where the coefficient obtained for endoparasitoid species is higher than for ectoparasitoid species. All statistical summaries of the Bayesian GLM models can be found on the GitHub repository under the name : Lifestyle_statistical_analysis_results.xlsx.

To further test the apparent correlation between Hymenoptera lifestyle and the rate of endogenization events, we inferred ancestral lifestyles along the phylogeny using a Bayesian model (see details in Materials and methods). We then constructed a generalized linear model where the dependent variable is the number of endogenization events inferred on each branch, while branch length and lifestyle are the explanatory variables (see details in Materials and methods). Branch length was included as an additive effect to remove the expected effect of time on the number of endogenization events, thus allowing the decomposition of the remaining variance according to the lifestyle (free-living, ectoparasitoid or endoparasitoid).

We first tested whether the rate of endogenization events deriving from any virus (that is, regardless of their genomic structures) was structured by lifestyles, and found no significant effect (Figure S5-A left side). We then split the dataset according to the genomic structure of the donor viruses. For RNA or ssDNA virus, the analysis did not reveal evidence of a correlation between wasps’ lifestyles and the rate of endogenization events (for details, see supplemental information, FigureS5-G,I K). On the contrary, in the case of dsDNA viruses, we found a highly significant effect of the wasp lifestyle: endogenization rates appear to be 2.47 times higher in endoparasitoids than in free-living species (89% CI [1.56-3.56], Figure4-A). The corresponding probability of direction (pd, an index representing the confidence in the direction of an effect) was equal to 99.9%. In contrast, ectoparasitoids did not differ from free-living species (Figure4-A). Accordingly, more than 98% of the MCMC iterations led to a higher coefficient value for endoparasitoids than for ectoparasitoids (so-called *P_MCMC_* in Figure4-A). This effect was consistently found using high confidence scaffolds only (A-ranked scaffolds, FigureS5-C right side). We also carried out the same analysis without the 4 domestication cases previously mentioned in the literature (because including them in our data set could have skewed the results) and reached the same conclusion (FigureS5-E right and left sides). Overall, these results show that dsDNA viruses are more often endogenized in endoparasitoids than in free-living and ectoparasitoid species.

### Domestications of dsDNA viruses are most prevalent in endoparasitoid species

We then investigated whether lifestyles may explain the abundance of domestication events. A simple Fisher’s exact test approach revealed an enrichment in endoparasitoid species of domestication events involving dsDNA viruses (Benjamini-Hochberg adjusted p-value = 1.8e-03), whereas no deviation from the null hypothesis was detected for the other viral genomic structures (Figure 3-B).

We built upon the generalized linear models described above, in a Bayesian framework, to test whether lifestyle could also be a factor explaining the propensity of Hymenoptera to domesticate (and not simply endogenize) viral genes (see details in Materials and methods). We found that, domestication of dsDNA viruses are 3.68 times more abundant in endoparasitoids than in with free-living species (89% CI [1.72-6.17], pd =99.9%, Figure4-B). This effect was also detected when only high confidence candidates were considered (FigureS5-D right side), or if we removed the four known cases of domestication (FigureS5-F left and right side). In other viral categories, no convincing effect of the wasp lifestyle was detected (all pd<99%) (Figure S5-H J) except for a higher rate of domestication of ssRNA viruses in free-living than in ectoparasitoid species (Figure S5-L)

Two non mutually exclusive hypotheses may explain the high frequency of dsDNA virus domestication in endoparasitoids. First, it may simply stem from the higher rate of endogenization outlined above: a higher rate of entry would overall translate into a higher rate of domestication. Second, it may result more specifically from differences in the rate at which viral elements are domesticated after being endogenized. To disentangle these hypotheses, we built a binomial logistic regression model in a Bayesian framework, focusing on events involving dsDNA viruses, and specifying the number of domesticated events *relative* to the total number of endogenization events inferred. By controlling for the endogenization input (the denominator), these binomial models make it possible to test whether the probability of domestication after endogenization of dsDNA viruses is correlated with the lifestyle.

Based on this analysis, the probability that an endogenization event will lead to a domestication event is not significantly different between endoparasitoids and freeliving species (FigureS10-A, pd=89.18%). However, the probability of domestication was found to be significantly higher in endoparasitoids than in ectoparasitoids (FigureS10-A, *P_MCMC_* =99.81%). The same trend was observed if we focused on high confidence scaffolds and/or if we removed the 4 known controls from the dataset (FigureS10-B,C D, pd < 86%).

Together, these findings show that the endoparasitoid lifestyle is associated with an increased rate of dsDNA viruses endogenization. Endoparasitoids are also characterized by an elevated frequency of domestication events that does not appear to be explained by an elevated rate of post-endogenization domestication.

### New remarkable cases of endogenization and domestication

Here, we describe in more details specific cases identified by our pipeline. We found a massive entry of genes from dsDNA viruses in an undescribed species belonging to the Campopleginae subfamily (”Campopleginae sp” in Figure 1). In Ophioniformes (a clade that includes Campopleginae), two lineages have previously been shown to host domesticated viruses (the Campopleginae species *Hyposoter didymator* [29], and the Banchinae species *Glypta fumiferanae* [30]). It has been advocated that these so-called ichnoviruses found in *Hyposoter didymator* and *Glypta fumiferanae* may derive from the same endogenization event [30]. In our unknown Campopleginae species, we identified homologs of 35 out of the 40 ichnovirus genes present in the genome of *H. didymator* (so-called IVSPER genes, [16]). Those genes show conserved synteny in the two species (Figure S11), strongly suggesting that they derive from the same endogenization event. However, our analysis did not identify viral homologs in the two Ophioninae and Cremastinae sufamilies, that are internal to the clade including Campopleginae and Banchinae wasps. This result argues against the view of a single event at the root of Ophioniformes, and thus supports the alternative view [31] that the so-called IVSPER genes in the Campopleginae and Banchinae subfamilies stem from independent events, despite their striking structural similarities (see FigureS12 for illustration). We found no trace of the previously suggested remnants of ichnoviruses in the related species *Venturia canescens*, whereas the presence of nudiviral genes in this species was confirmed [17].

We found 5 new cases of endogenization involving multiple EVEs from dsDNA viruses belonging to *Nudiviridae*, LbFV-like and AmFV-like families.

Two of them involve parasitoid species, i.e. *Platygaster orseoliae* and an *Aprostocetus* species. For *Aprostocetus*, we detected 6 EVEs related to nudiviruses branching between the Chalcidoidea and the Diaprioidea superfamilies (Fig. S4). Among these EVEs we found four with an annotation : *lef-4*, *Ac68/pif-6*, GrBNV_gp19/60/61-like proteins, and a rep-like protein. In the absence of closely related sequences or RNA seq data, we cannot investigate if these elements have been domesticated. The *P. orseoliae* case involve the recently characterized putative family of filamentous viruses [32]. The free-living LbFV virus is the only representative of this putative family and has been identified as a source of adaptive genes in *Leptopilina* wasps that parasitize *Drosophila* flies, with 13 virally-derived genes involved in the production of VLPs protecting the wasp’s eggs from encapsulation [19]. In *P. orseoliae*, 15 genes homologous to LbFV were detected (out of 108 ORFs in the LbFV genome; median E-value = 9.39e-21 [min = 2.617e-76, max = 4.225e-08], Figure S8-A). Among these 15 genes, 5 were also endogenized in *Leptopilina* species (named LbFV_ORF58:DNApol, LbFV_ORF78, LbFV_ORF60:LCAT, LbFV_ORF107 LbFV_ORF85) [19]. Assuming the ancestral donor virus contained the same 108 genes as LbFV, the number of shared genes in these two independent domestication events is higher than expected by chance (one-sided binomial test: x = 5, n = 15, p = 13/108, p= 0.02682), suggesting that similar functions could have been retained in both lineages (see additional information in supplemental information). Notably, we also found within the *P. orseoliae* assembly a new “free-living” virus among scaffolds noted as F or X, related to LbFV, that we propose to call PoFV (Platygaster orseoliae filamentous virus) (see fig. 5 and supplemental information for details). This virus is the closest relative to the EVEs found in *P. orseoliae* and is composed of 136 ORFs FigureS8-B). Using this putative whole genome viral sequence to search for homologous genes in the *P. orseoliae* genome, we were able to detect a total of 139 convincing EVEs (corresponding to 89/136 PoFV ORFs). 44/89 of these EVEs presented signs of domestication using paralogs, with dN/dS significantly lower than 1 (see details in supplemental information). Although functional studies are clearly needed to confirm that these virus-derived genes are involved in the production of VLPs as in *Leptopilina* [19], we see Platygaster orseoliae endogenous viral elements (PoEFVs) as good candidates for viral domestication, which could possibly be involved in counteracting the immune system of its dipteran host (from the Cecidomyiidae family [33]). To our knowledge, this is the first report of a massive viral endogenization and putative domestication within the Platygastroidea superfamily.

**Figure 5.**
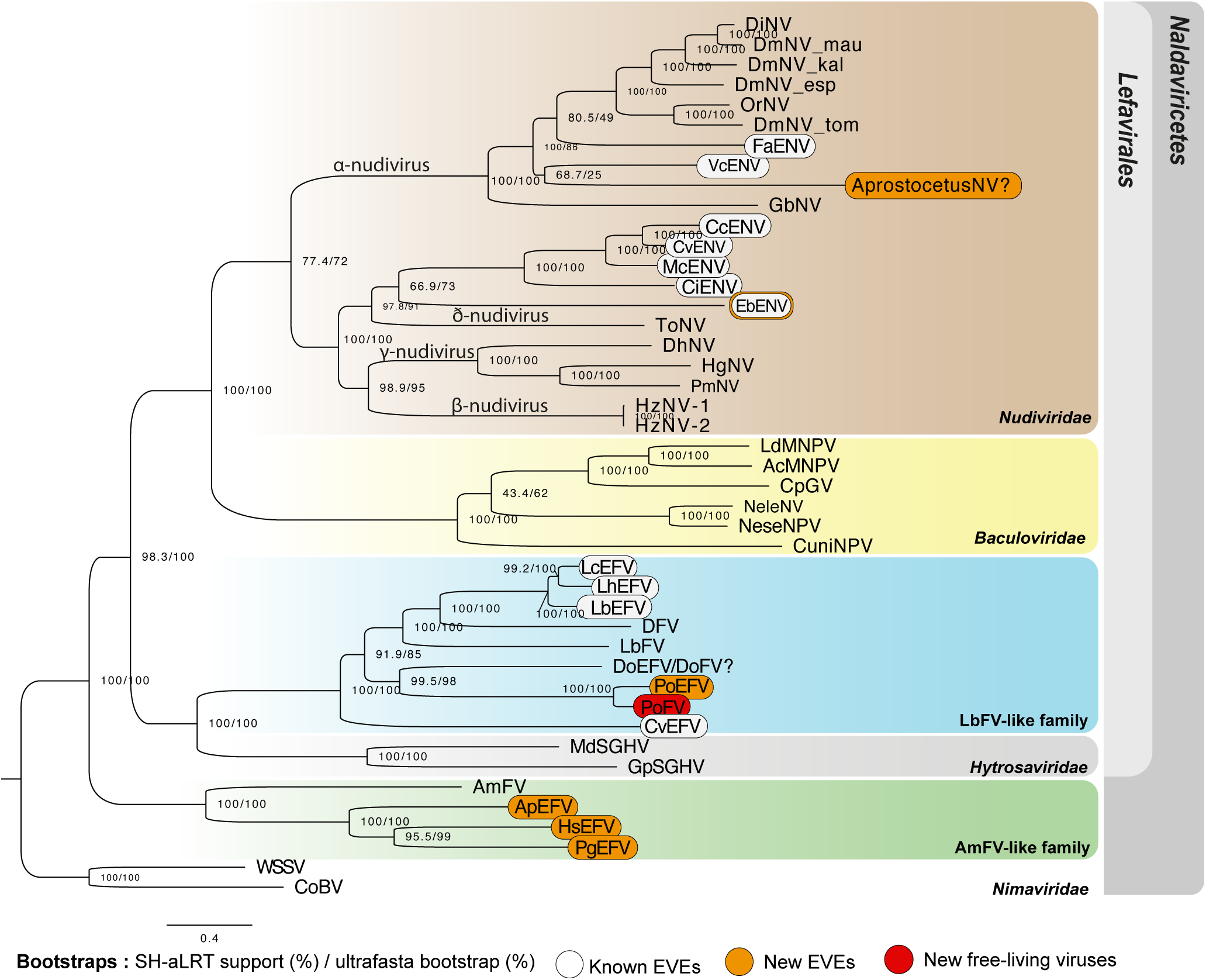
Phylogenetic relationships among endogenized and “free-living” dsDNA viruses. Specifically, this figure shows the relationships between *Naldaviricetes* double-stranded DNA viruses and EVEs from hymenopteran species, where at least 3 endogenization events were found. This tree was computed using maximum likelihood in Iqtree (v2) from a 38,293 long protein alignment based on the concatenation of 142 viral genes. Confidence scores (aLRT%/ultra-bootstrap support%) are shown at each node. The scale bar indicates the average number of amino acid substitutions per site. Previously known EVEs are in white, those from the present study in orange, and leaves inferred as free-living viruses are in red. All the best partitioned models can be found on the Github repository under the name : dsDNA_phylogeny_best_ML_partitions.nxs. All free-living dsDNA viruses used in this phylogeny were obtained from published complete viral genomes. More details on the phylogenetic inference can be found in methods.

The other three cases involved ant species : *Harpegnathos saltator* (EsEFV) (12EVEs/6dEVEs), *Pseudomyrmex gracilis* (PgEFV) (9EVEs/1dEVE), *Aphaenogaster picea* (ApEFV) (7EVEs). These endogenized elements are related to a poorly characterized family of filamentous viruses denoted AmFV [34, 35]. In *H. saltator*, 9 genes deriving from an AmFV-like virus were detected (including 3 genes that have been previously identified by [23]). Intriguingly, all these genes presented numerous paralogs within the genomes (135 in total) (FigureS7 FigureS4), with 22 copies for AmFV_0062 (*pif-1*), 18 for AmFV_0102 (*pif-2*), 51 for AmFV_0090 (*pif-3*), 24 for AmFV_0044 (*integrase*), 13 for AmFV_0079 (*p74*), 5 for AmFV_0047 (*RNA polymerase*), 19 for AmFV_0126 (Unknown), 23 for AmFV_0168 (Unknown) and 7 for AmFV_0154 (Unknown). Most paralogs were found in scaffolds exceeding the expected size for any virus sequence (min = 23,726bp, mean = 326,262bp, max = 2,693,376bp). In addition, all scaffolds do include transposable elements and eukaryotic genes making them undoubtedly endogenized. Accordingly, our pipeline attributed the highest confidence index A for 104 of them (out of 135). The *P.gracilis* genome revealed 9 EVEs, including homologs of *pif-1*, *pif-3*, *RNA polymerase*, *ac81*, *integrase* and *odv-e56* (FigureS4). Notably, one of the 9 EVEs (AmFV_059, of unknown function) shows both a dN/dS < 1 (mean= 0.1747, p-value 5.877e-02), and a very high TPM value (362836 TPM from whole body tissues). Finally, in *A. picea*, 7 EVEs were detected, including homologs of *pif-1*, *pif-3*, *integrase*, *odv-e56* and *p74* (Figure S7). No raw reads data were available for this species, precluding coverage-based inferences. Since there were neither orthologs or paralogs for these genes to compute dN/dS analyses, nor transcriptomic data, it was not possible to infer their domestication status. At this stage it is thus not possible to conclude as to the functions of these genes in *H. saltator*, *P. gracilis* and *A. picea*, but this surely deserves further attention.

## Discussion

All kinds of viruses can integrate arthropod genomes, although the mechanisms underlying these phenomena remain unclear [1, 4]. Prior to the present analysis, 28 viral families had been described as involved in endogenization in arthropods [4]. Our study of Hymenopteran genomes further revealed the ubiquity of this phenomenon, with at least 40 viral families (or family-like clades) involved. Of the 1,261 EVEs found, the average identity with the closest known viral proteins was 36.32% [min = 15.7%, max = 99.1%]. Although this large overall divergence suggests ancient events, it does not exclude the possibility that some of the integrations are recent, because free-living viral relatives of the true donors may be unknown or extinct [36].

In the following section, we will first discuss why double-stranded DNA viruses, in comparisons with other viral genomic structures (ssDNA, dsRNA, ssRNA), are more often endogenized than expected. We will then discuss hypotheses that could explain why we found a higher rate of endogenization of dsDNA viruses among endoparasitoids compared to ectoparasitoids or free-living hymenopterans, which also translates into more frequent events of domestications.

### dsDNA viruses are more frequently involved in endogenization than expected by chance

Despite the observations that all viral genomic structures can be involved in endogenization, we clearly identified differences in their propensity to do so. Based on a comparison between the respective proportions of the various viral categories in the inferred endogenization events and in public databases, we found that dsDNA viruses are much more represented than expected, while ssRNA viruses are under-represented (Figure 2-A). We acknowledge that current knowledge on the actual diversity of free-living viruses (as approximated through the NCBI taxonomy database) remains incomplete, but the strength of the effect reported here makes this conclusion rather robust to variations in the null distribution. On the basis of current knowledge, RNA viruses, and in particular ssRNA viruses, appear to be much more diversified and prevalent than DNA viruses in insects. We note that viral-metagenomic studies often focus either on DNA or RNA viruses, and as such do not provide an accurate and unbiased picture of the extent viral diversity. To gain insights on this topic, we may thus focus on model systems where long-lasting research efforts have likely produced a more reliable picture. The Honeybee *Apis mellifera* is probably the most studied of all Hymenopteran species. In honeybees, the great majority of known viruses belongs to the RNA world [37], with very few exceptions [35]. Similarly, until 2015, only RNA viruses were known to infect the fruit fly *Drosophila melanogaster*, despite the extensive research conducted on this model system [38]. A very limited set of DNA viruses has now been described from this species [39] but clearly, RNA viruses dominate the *Drosophila* viral community, both in terms of diversity and prevalence. In support of this view, recent studies revealed the very elevated absolute diversity of RNA viruses. For instance, a survey of 600 insect transcriptomes recovered more than 1,213 RNA viruses belonging to 40 different families [40]. Although, obviously, this study does not inform on the diversity of DNA viruses, it shows that the RNA virome of insects is both prevalent (e.g., in this study, 15% of all insects were infected by a single Mononegales-like virus) and extremely diversified [40]. Actually, this view appears to hold at the larger scale of eukaryotes [41]. Taking into account this patent abundance of ssRNA viruses in insects, our study indicates they are by far less frequently endogenized than their dsDNA counterparts in hymenopterans. Notably, a similar trend was recently reported in a study including a diverse set of eukaryotes [24].

Most of the major endogenization events characterized so far in hymenopterans involve dsDNA viruses from the *Nudiviridae* family [21, 18, 17, 42, 43, 44, 4]. Our study further confirms that this viral family represents a major source of exogenous and sometimes adaptive genes for Hymenoptera. Indeed, 28 new independent endogenization events involve this family, among which 9 are shared by at least two related species (Figure 2-B, Figure1). The major contribution of nudiviruses to endogenization may be explained by their wide host range in arthropods [45]. Their nuclear replication constitutes another plausible explanatory factor [46], since it may facilitate contact with host DNA. In addition, their tropism for gonads may favor the endogenization in germinal cells [47]. In fact, nuclear replication is a feature shared by nearly all families of dsDNA viruses found in our analysis : *Baculoviridae*, *Iridoviridae*, *Phycodnaviridae*, *Nimaviridae Caulimoviridae*, *Herpesviridae*, *Asfaviridae* (at early times) [48, 49, 50, 51, 52], Apis-filamentous-like [53] and LbFV-like families [54] (the *Poxviridae* viruses, that replicate in the cytoplasm, are thus the only exception). In contrast, most RNA viruses replicate in the cytoplasm. Nuclear replication may thus constitute a general explanation for the elevated propensity of DNA viruses to endogenization. Additionally, we may expect that a DNA molecule, rather than an RNA molecule, is more likely to integrate the insect genome, because the latter requires reverse transcription before possible endogenization.

The *Poxviridae* case indicates that cytoplasmic replication does not necessarily impede endogenization. These viruses do not require nuclear localization to propagate [55, 48] and were nevertheless found to be involved in many endogenization events (n=28). A similar pattern was observed in a recent study focusing on ant genomes [23]). Within Poxviridae, entomopoxviruses were particularly involved in endogenization events (n=18) with four cases of EVEs shared between several closely related species (Figure 2-B).

### Factors behind variations in endogenization and domestication rates

Several recent studies have uncovered abundant EVEs in insect genomes [23, 20, 56], with huge variation in abundance between species. For instance, in their analysis based on 48 arthropod genomes, [20] found that the number of EVEs ranged between 0 and 502. Although insect genome size and assembly quality may partly explain this variation [4], the underlying biological factors are generally unknown. In this study, we tested the hypothesis that the insect lifestyle may influence both the endogenization and domestication rates. We used a Bayesian approach to reconstruct ancestral states throughout the phylogeny of Hymenoptera, thus accounting for uncertainty, and found that endoparasitoidism, in comparison with other lifestyles, tends to promote dsDNA viral endogenization. Notably, this conclusion was not the artefactual consequence of differences in genome assembly quality. In fact, the quality of genome assemblies was correlated with the lifestyle in our data set, but the genomes of endoparasitoid species were generally less well assembled than those of free-living species. If anything, this difference should reduce the power for detecting endogenization events in endoparasitoids, where our analysis detected an excess of such events. Our estimate of the effect sizes (with 2.47 times more endogenization events in endoparasitoids than in free-living species) should thus be seen as conservative. Why do endoparasitoid wasps tend to undergo more endogenization than others? We initially had in mind two non-exclusive hypotheses that remain plausible explanations for the observed pattern. First, endoparasitoids may be more intensively exposed to viruses. In addition, or alternatively, endoparasitoids may have a higher propensity to endogenize and retain viral genes.

Several factors come in support of the first hypothesis. Endoparasitoid larvae grow by definition inside their host’s body, and such a close interaction implies that any endoparasitoid individual will also be interacting with its host’s viruses. Accordingly, the best studied cases of viral domestication in wasps involve nudiviruses, that are known to replicate in their caterpillar hosts [57, 17, 42]. Another putatively important factor is the presence of virus in the venoms that parasitoids inject into hosts together with their eggs. These are known to protect the offspring against the host’s immune response, and to manipulate the host physiology [58] but this feature could favor the subsequent spread of viruses in wasp populations: by colonizing the venom-producing tissues (venom gland or calyx, depending on the species biology) viruses may secure an effective pseudo-vertical transmission and thus maintain themselves efficiently in wasp populations. Numerous endoparasitoid viruses benefit from such pseudo-vertical transmission [54, 59], including some whose relatives have been endogenized and domesticated by endoparasitoids [19]. The presence of viruses within venoms may also facilitate horizontal transmission between conspecifics in the case of superparasitism, as observed in the *Drosophila* parasitoid *Leptopilina boulardi* [27]. Although this effect may also be at play for some ecto-parasitoid species [60], we expect it to be more pronounced for endo-parasitoid species since they have a closer interaction from the inside of their hosts. Generally, endoparasitoids may thus carry a higher load of non-integrated viruses than other hymenopterans. However, if this effect is at play, we expect to have an “endoparasitoid” effect for all viruses, whatever their genomic structure. For instance, we would expect such an effect to be detected for ssRNA viruses, which are involved in the greatest number of endogenization events (Figure2-A). This was not the case, since only dsDNA viruses were more frequently endogenized in endoparasitoids. Thus, we argue that this hypothesis is unlikely to explain the observed pattern.

The second hypothesis posits that endoparasitoids are more frequently selected for retaining virally-derived genes than ectoparasitoid or free-living hymenopterans. In our analysis, domestication events are most frequently observed in endoparasitoids (over 3 times more frequently than in other hymenopterans). Obviously, this may be at least partly explained by the higher input discussed above (the higher endogenization rate). Yet, once this effect is controlled for, a trend towards a higher rate of domestication remains. More specifically, the likelihood of domestication following endogenization was significantly higher in endoparasitoids than in ectoparasitoids, but was not significantly higher than in free-living species. This latter lack of significant difference may be biologically explained if a single domestication event precludes the domestication of additional EVEs, while not affecting the rate of non-adaptive endogenization. This would “dilute” the signal along branches involved in domestication. If this effect is at play, then it reduces considerably the power of our analysis to detect any difference on the rate of domestication between lifestyles. Indeed, in all known cases, only one domesticated virus has been documented, suggesting that further domestications are not beneficial once a viral machinery has been recruited by a wasp lineage.

Whether or not the rate of domestication *per se* is higher in endoparasitoids than in other hymenopterans, the selective advantages brought by these viral genes in endoparasitoids should be discussed. It has been demonstrated in a few model systems that EVEs may confer antiviral immunity against related “free-living” viruses via the piRNA pathway [7, 61]. Yet, to our knowledge, such an effect has only been demonstrated against RNA viruses, so that it would not explain the excess of DNA viruses documented here. Furthermore, the sequence identities with known viral sequences, which is needed for this mechanism to work, is low in our dataset. Accordingly, previous work revealed that EVE-derived piRNAs studied in 48 arthropod species were also probably too divergent to induce an efficient antiviral response [20]. At that stage, the ability of EVEs to generate PIWI-interacting RNAs that play a functional role in antiviral immunity seems questionable. Further studies involving small RNA sequencing in hymenopterans would be required to shed light on this issue. Protection of the eggs and larvae against the host immune system is recognized as an important trait, where EVEs play a critical role. Because of their peculiar lifestyle, endoparasitoids are all targeted by the host immune system, a matter of life or death to which other hymenopterans are not exposed to. Several cases of endogenization and domestication in endoparasitoids, all involving dsDNA viruses, are thought to be related to this particular selective pressure [42, 16, 17, 18, 19]). The parasitoids appear to have co-opted the viral fusogenic property to address their own proteins (VLPs) or DNA fragments (polydnaviruses) to host immune cells, thereby canceling the host cellular immune response. The above-hypothesized high exposure of endoparasitoids to viruses, together with this unique selective pressure, may act in concert to produce the pattern documented here: a strong input, that is, a diverse set of putative genetic novelties, combined with a strong selective pressure for retaining some of them. The observed excess of dsDNA viruses may be an indication that these viruses display a better potential for providing adaptive material in this context. In the cases of polydnaviruses (found in some Braconidae and some Campopleginae), it appears that one way to efficiently deliver virulence factors to the host cell is by addressing DNA circles that ultimately integrate into the host immune cells and get expressed [62, 63]. The DNA which is packed into the mature particles typically encodes virulence proteins deriving from the wasp [64]. This means that, at least for these cases, the viral system should be able to pack DNA, which is most likely a feature that DNA viruses may provide. Such an argument does not hold in the VLP systems, where only proteins are packed in viral particles, and it is unclear why EVEs deriving from dsDNA viruses would be more able to fulfill such a function. Here other features of dsDNA viruses come into mind as possibly important factors: their large genome size, and their large capsids and envelopes [65]. These may predispose dsDNA viruses to be domesticated, since abundant quantities of venoms have to be transmitted in order to efficiently suppress the host immune response.

## Conclusion

Our analysis has revealed a large set of new virally-derived genes in Hymenoptera genomes. Those genes were deriving from viruses with any genomic structures, although dsDNA viruses were disproportionately involved in endogenization and domestication. Importantly, our analysis revealed that endogenization rate and the abolute number of domestication events involving dsDNA viruses was increased for endoparasitoids compared to other lifestyles. Among the new cases of endogenization and domestication, we uncovered new events revealing common features with previously known cases of viral domestication by endoparasitoids, such as in the Platygastroidea *Platygaster orseoliae*. This is to our knowledge the first case reported in the superfamily Platygastroideae, thus extending the diversity of Hymenoptera concerned by viral domestication. We propose that the higher rate of endogenization and higher number of domestication events in endoparasitoids is a consequence of the extreme selective pressure exerted by the host immune system on endoparasitoids. This extreme selective pressure may select endoparasitoids for retaining a viral machinery that could help them address virulence factors to their hosts. We expect this process to be widespread among insect species sharing the same lifestyle.

## Materials and methods

### Genome sampling, assembly correction and assembly quality

A bioinformatic pipeline mixing sequence homology search, phylogeny, genomic environment, and selective pressure analysis was built to search for viral endogenization and domestication events in Hymenoptera genomes. We used 133 genome assemblies in total, of which 101 were available on public repositories (NCBI and BIPPA databases) and 32 were produced by our laboratory (all SRA reads and assemblies available under the NCBI submission ID : SUB11373855). Concerning the last 32 samples, DNA was extracted on single individuals (usually one female) or a mix of individuals when the specimens were too small using Macherey-Nagel extraction kit, the DNA was then used to construct a true seq nano Illumina library at Genotoul platform (Toulouse, France). The sequences were generated from HiSeq 2500 or HiSeq 3000 machines (15Gb/sample). The paired-end reads were then quality trimmed using fastqmcf (-q15 –qual-mean 30 -D150, GitHub) and assembled using IDBA-UD [66]. All sample information can be found on the GitHub repository under the name : All_sample_informations.txt and is available under the NCBI Biosample number : SUB11338872.

The size of the 133 assemblies ranged from 106.14mb to 2102.30mb. We kept only genome assemblies containing at least 70% non-missing BUSCO genes (124/133 genomes, [67]) (all genome information can be found on the GitHub repository under the name : Assembly_genome_informations.txt). In addition, when the raw reads were available, we used the MEC pipeline [68] to correct possible assembly errors. Although some genomes were highly fragmented (such as the 32 genomes we generated since they were obtained using short reads only), the N50 values (min: 3542bp) were equal to or larger than the expected sizes of genes known to be endogenized and domesticated (min known domesticated EVE : 165bp) indicating that most of the putative EVEs should be detected entirely.

Out of the 32 samples sequenced by our laboratory for this study, one (corresponding to *Platygaster orseoliae*) gave unexpected results. After assembly and BUSCO analysis, two sets of contigs were identified: one with only 4X coverage on average, and one with 33X on average. The phylogeny of these different BUSCOs gene sets showed that the low-coverage scaffolds likely belong to an early diverging lineage of Chalcidoidea (Figure1), whereas the 33x scaffolds belong to the target species *P. orseoliae*. This result suggests that the pool of 10 individuals used for sequencing was likely a mix of two species. A phylogenetic study based on Ultra Conserved Elements (UCEs) obtained from several species of Chalcidoidea [69, 70] allowed to identify the unknown species as sister to *Aprostocetus sp* (Eulophidae) (see details in the supporting information and Figure S2). In the figures and tables, the name putative_ *Aprostocetus*_sp was consequently assigned to the unknown sample. However, since the lifestyle and identity of this species are uncertain, we did not include the corresponding scaffolds in the main analysis. The scaffolds belonging to this putative_*Aprostocetus*_sp. (i.e : all scaffolds with a mean coverage < 10X) were removed from the *P. orseoliae* assembly file hosted in NCBI.

### Pipeline outline

EVEs were identified from the 124 Hymenoptera assemblies using a sequence-homology approach against a comprehensive viral protein database (including all categories of viruses : ssDNA, dsDNA, dsRNA and ssRNA). In order to validate viral endogenization within Hymenoptera genomes, we developed an “endogenization confidence index” ranging from A to X (FigureS13-7). This index takes into consideration the presence of eukaryotic genes and/or transposable elements around candidate loci, and scaffolds coverage information (coverage for a valid candidate should be similar to that found in BUSCO containing scaffolds). Finally, the pipeline also included an assessment of the evolutionary history and of the selective regime shaping the candidates (based on dN/dS and/or expression data).

### Hymenoptera phylogeny

The phylogenetic reconstruction of the 124 Hymenoptera species was performed based on a concatenation of the 375 BUSCO proteins. The analysis was conducted by maximum likelihood via Iqtree2 [71] selecting the best model [72]. The tree was rooted via two species of the Coleoptera order (*Anoplophora glabripennis* and *Tribolium castaneum*). Bootstrap scores were evaluated using the UFboot approach [73]. The results found were consistent with a previous, more comprehensive study [28].

### Search for viral homology

We collected all protein sequences available in NCBI virus database [74], removing phage and polydnavirus (virulence genes from wasp origin found within PDVs) sequences. This database contained 849,970 viral protein sequences (download date : 10/10/2019), to which the 40 putative viral proteins encoded by the *Hyposoter didymator* genome were added (so-called IVSPER sequences, [29]). The sequence homology search was performed with a BlastX equivalent implemented in Mmseqs2 [75]) using each genome assembly as queries and the viral proteins collected as database. The result gave a total of 81,953,678 viral hits (max E-value 5e-04 with an average of 660,916 hits per genomes). We kept only candidates with a percentage coverage of the viral protein >= 30%, an identity score >= 20% and an E-value score < 5e-04 (Figure S13-1). The threshold parameters were optimized to maximize the detection of the 13 endogenous viral sequences within the genus *Leptopilina* [19]. Once all the viral hits were recovered, we formed putative EVEs loci (n=238,108) corresponding to the overlap of several viral hits on the same scaffold using the GenomicRanges R package [76] (Figure S13-2). To remove false positives corresponding to eukaryotic genes rather than viral genes, we then performed another generalist sequence homology search against the Nr database (downloaded the 09/11/20) using mmseqs2 search (-s 7.5, E-value max = 0.0001) (Figure S13-3). We did not select our candidate based on the best hit, since it does not necessarily reflect the true phylogenetic proximity. Instead, candidates with more than 25 hits with either eukaryotic non-hymenoptera species or prokaryotic species were removed, except if they also had hits with at least 10 different virus species (bits >= 50). We chose to eliminate Hymenoptera hits from the database because if a real endogenization event concerns both one of the 124 species of our dataset and some species in the NCBI database, then an apparent “Hymenoptera” hit will be detected, possibly leading to its (unfair) elimination. Since viral diversity is poorly known, we also kept sequences with even one single viral hit, as long as it did not have more than 5 eukaryotic or prokaryotic hits. Using these filtering criteria we removed a total of 234,036/238,108 (98,3%) candidate loci leaving 4,072 candidates with convincing homology to viral proteins. Note that among these loci a certain proportion actually corresponded to non-endogenized “free-living” viruses. To study the evolutionary history of these candidate EVEs, we then performed a general protein clustering of all the candidates and the NCBI viral proteins (Figure S13-4, Mmseqs cluster; thresholds : E-value 0.0001, cov% 30, options : –cluster-mode 1 –cov-mode 0 –cluster-reassign –single-step-clustering[77]).

We eliminated from the dataset the chuviral glycoproteins that have been captured by LTR retrotransposons [78], as these loci have complex histories mostly linked to the transposition activity after endogenization. For this purpose, we systematically searched among the candidates for the presence of TEs within or overlapping with the EVE (see the file All_TEs_overlapping_with_EVEs under the github repository). Only one cluster (Cluster4185) was concerned by such a situation (chuviral glycoproteins overlapping to Gypsy/LTRs). It was detected in 89/124 species (1074 total copies, median = 7 copies/species, max = 244, min =1), and was probably similar as the one described in [79].

### Evolutionary history and selection pressure acting on endogenous loci

#### Arguments for endogenization

Among all the candidates for endogenization there were probably false positives that corresponded either to natural contaminants (infecting viruses sequenced at the same time as the eukaryotic genome) or laboratory contaminants (virus accidentally added to the samples). One way to filter these cases was to study (i) the genomic environment (are there other eukaryotic genes or transposable element on the same scaffolds?) and (ii) metrics such as G+C% (used only for read coverage/GC plots) and scaffold coverage depth around candidate loci (are they the same as scaffolds containing housekeeping genes?). All of these data were used to establish confidence in the endogenization hypothesis, scaled from A to X (Figure S13-7).

**(i) Scaffolds sequencing depth (Figure S13-5) :** In order to support the hypothesis that a scaffold containing candidate EVEs was part of the Hymenoptera genome, we studied the sequencing depth of the scaffolds. If the sequencing depth of a candidate scaffold was not different from the depth observed in scaffolds containing BUSCO genes, then this scaffold was likely endogenized into the Hymenoptera genome. Hence, when DNA reads were available (FigureS1), we measured this metric by mapping the reads on the assemblies using hisat2 v 2.2.0 [80]. An empirical p-value was then calculated for each scaffold containing a candidate EVE. To calculate this empirical p-value, we sampled 500 loci of the size of the scaffold of interest within BUSCO scaffolds. These 500 samples represented a null distribution for a scaffold belonging to the Hymenoptera genome. The p-value then corresponded to the proportion of BUSCO depth values that were more extreme than the one observed in the candidate scaffold (two-sided test). We used a threshold of 5% and a 5% FDR (multipy python package [81]).

**(i) Genomic environment and scaffold size** (Figure S13-6) Another way to rule out contaminating scaffolds was to look for the presence of eukaryotic genes and transposable elements in the scaffolds containing candidate EVEs, assuming that their presence in a viral scaffold is unlikely. Indeed, so far, very few viral genomes have been shown to contain transportable elements [82, 83, 84, 85, 86] because TE insertions are mostly deleterious and are therefore quickly eliminated by negative selection [84, 85]. We searched for transposable elements by a BlastX-like approach (implemented in Mmseqs2 search -s 7.5), taking as query the scaffolds of interest and as database the protein sequences of the transposable element (TE) available in RepeatModeler database (RepeatPeps, v2.0.2) [87]. We only kept hits with an E-value < 1e-10 and with a query alignment greater than 100aa. We then merged all overlapping hits and counted the number of TEs for each scaffold. To find eukaryotic genes within genomes we used Augustus 3.3.3 [88] with BUSCO training and then assigned a taxonomy to these genes via sequence homology with Uniprot/Swissprot database using mmseqs2 search [75], and only retained genes assigned to insects.

Accordingly, the scaffolds were scored as follows (Figure S13-7):

- A: scaffolds with a corrected coverage p-value > 0.05 and at least one eukaryotic gene and/or one repeat element,
- B: scaffolds with at least one eukaryotic gene and/or one repeat element but no coverage data available,
- C: scaffolds with a corrected coverage p-value > 0.05 and neither eukaryotic gene nor transposable element,
- D: scaffolds with a corrected coverage p-value < 0. 05 and whose coverage value was higher than the average of the scaffolds containing BUSCOs (as it is difficult to imagine that an endogenized scaffold present a lower coverage than expected, whereas a higher coverage could correspond to the presence of repeated elements that inflate the coverage of the scaffold for example) but with at least 5 eukaryotic genes and/or a repeated element (in total),
- E: scaffolds presenting no DNA seq coverage data available and no eukaryotic gene nor transposable element detected,
- F: scaffolds presenting a corrected p-value of coverage < 0.05 and less than 5 eukaryotic genes without any transposable elements; this category may rather correspond to free-living viruses.
- X: scaffolds with a corrected p-value < 0.05 and neither eukaryotic gene nor transposable element; This category may rather correspond to free-living viruses.

#### Inference of endogenization events

Because several EVEs may derive from the same endogenization event, we sought to aggregate EVEs into unique events. We aggregated into a single event, firstly (i) all the EVEs present on the same scaffolds, and secondly (ii) all the EVEs that presented the same taxonomic assignment at the level of the viral family. These two steps were sufficient to aggregate EVEs in the simplest case of events involving only one species (but possibly several EVEs).

To further characterize the endogenization events including more than one Hymenoptera species, we also relied on phylogenetic inference. To this end, the protein sequences belonging to each of the clusters (containing both viral proteins and candidate EVEs) were first aligned with clustalo v1.2.4 [89] in order to merge possible candidate loci (which may in fact correspond to various HSPs). All loci (=HSPs) within the same scaffold presenting no overlap in the alignment were thus merged, as they probably correspond to multiple HSPs and not duplications. We then performed a new codon alignment from the augmented sequences in the clusters using the MACSE v2 alignsequence program [90] (Figure S13-8). This alignment allowed us to obtain a protein and nucleotide codon alignment. We used the protein alignment to infer the phylogeny of each cluster with the program Iqtree2 v2.1.2 [71] (-m MFP -alrt 1000 (partitioned))(Figure S13-9). No trimming was performed at the amino-acid level, since this may result in loss of topology information [91, 92]. However, since it can affect branch length, only codon alignment was trimmed at the protein level via Trimal v1.2 (Figure S13-10) (- backtrans -automated1) [93]. We then exploited the information from the cluster phylogenies to form the endogenization events. EVEs potentially deriving from the same event should be supported by the formation of the same well-supported monophyletic clade (bootstrap > 80) both in the gene tree and the Hymenoptera tree (allowing gene losses in 20% of the species concerned by the monophyletic group). EVEs were possibly aggregated within the same event only if the Hymenoptera belonged to the same family. (Figure S13-11). Finally, the clustering of multiple EVEs within the same scaffold in one species was used to aggregate the homologous EVEs found in a related species within the same shared event, even if they were on different scaffolds (Figure S13). For details, see some canonical examples in Figure S14.

For events shared by several species, we were also able to analyze gene synteny around putative EVEs. To do this, we conducted the equivalent of an all vs all TblastX (Mmseqs2 search –search-type 4, max E-value =1e-07) between all the candidate loci within a putative event (deduced from the phylogenetic inference), and then looked for hits (HSPs) between homologous EVEs around the insertions. Because it is possible to find homology between two genomic regions that does not correspond to orthology, for example because of the presence of conserved domains, we had to define a threshold to identify with confidence the orthology signal. We therefore conducted simulations to define this value, based on the well assembled genome of *Cotesia congregata* (GCA_905319865.3) by simply performing the same all vs all blast analysis against itself (as if the two species considered had the same genome). Based on this, we defined two types of simulated EVEs, (i) independently endogenized EVEs in the genomes of the two “species”. This is simply simulated by randomly selecting two different regions in the genomes, and (ii) a shared simulated EVE that was acquired by their common “ancestor”. This is simulated by selecting the same random genomic location in both “genomes”. We then counted the total length of the HSPs found around the simulated insertions all along the corresponding scaffold (i and ii). As the result will obviously depend on scaffold length, we performed these simulations on several scaffold lengths (100000000bp, 10000000bp, 1000000bp, 100000bp and 10000bp). We conducted 500 simulations in each scenario, and we measured the cumulative length of homologous sequences by counting the sum of HSPs (bit score > 50). We then defined a threshold for each windows size in order to minimize for the false-positive (FP) and maximize true-positifs (TP) (thresholds 100000000bp = 172737bp (FP = 0.012, TP= 0.922); 10000000bp = 74262 bp (FP= 0.012, TP=0.878) ; 1000000bp = 21000 bp (FP=0.014, TP=0.28); 100000bp = 1332 bp (FP= 0.012 TP= 0.198) and 10000bp = 180 bp (FP= 0.008, TP= 0.208)).

Events were linked to viral families based on the closest match information between the viral blastx (GenBank accession number and/or viral protein and/or viral species) and the classification proposed in [94].

#### Arguments for domestication

One way to test for the domestication of an EVEs (dEVEs) was to estimate the ratio (omega) of the number of nonsynonymous substitutions per non-synonymous site (dN), to the number of synonymous substitutions per synonymous site (dS). If EVEs are evolving neutrally, then the ratio is expected to be equal to 1, whereas if the EVE is under purifying selection, dN/dS is expected to be lower than 1. We conducted this analysis on trimmed codon alignments from (Figure S13-11) via the codeml algorithm from PAML [95] used through the ETE3 package [96] (model Muse and Gaut [97]). We then used a branch model to test the deviation from the null model in which marked branches (called foreground) wich corresponded to the monophyletic EVE clade evolved under a neutral scenario (*x*^2^ test). The dN/dS estimated for the whole clade is then the average of each branch of the clade. The p-values were then adjusted by selecting an FDR of 0.05 [81], and we estimated the standard errors of dN/dS that maximized the likelihood (option getSE = 1). dN/dS with dS greater than 10 were removed, since this indicates substitution saturation (Figure S13-12).

The other way we choose to study the domesticated nature of a viral gene was to study their expression profile (Figure S13-13). We reasoned that domesticated genes are likely to be significantly expressed. To test this, when RNAseq reads were available on NCBI (SRA), we mapped them on the assembled genomes (until reaching 300x coverage as far as possible). Using the TPMCalculator program [98], we measured expression in ovaries and whole body if available or alternatively in any tissue (see supplemental information table :RNA_seq_reads_mapped.txt). An EVE was considered as domesticated if the gene was expressed with a Transcripts Per Kilobase Million (TPM) index above 1000. This threshold was chosen based on the median value observed for control EVEs (718.70 TPM), rounded up to 1000TPM to be conservative. We measured the accuracy of this metric using EVEs for which both TPM and dN/dS calculations were possible: among the 36 genes having a TPM>1000, 33 also had a dN/dS significantly below 1 suggesting that inferring domestication based on TPM>1000 was consistent with dN/dS test with a 0.9166 probability. Finally, based on the idea that an active EVE should encode a protein with similar length to the donor virus, we calculated the actual viral protein sequence length using the orfipy algorithm [99] (Figure S13-14).

A possible bias when comparing the effect of lifestyles on domesticated elements could come from a difference of RNAseq reads availability depending on the lifestyle, which may result in a different number of EVEs considered as domesticated. A GLM binomial analysis did not reveal any correlation between RNAseq data availability and lifestyle (endoparasitoid = Slope(SE)=0.21(0.62), p=0.73; free-living= Slope(SE)=0.40(0.57), p=0.49 using ectoparasitoid as intercept).

### Sensitivity and specificity of the analysis

#### Capacity to find Endogenous Viral Elements (EVEs)

Among the species included in our dataset, 7 were known to contain a domesticated virus (2 with similar PDV [42], 5 with different VLPs [17, 18, 19]), corresponding to 4 independent endogenization events. Our pipeline was able to detect the vast majority of the corresponding virally-derived genes (88.6%, details in table S1). The 11.14% false negatives corresponded to sequences that were too divergent or with a region of similarity too small to be detected by our pipeline. We found that 88.7% of the control EVEs were located within scaffolds scored as A (i.e. having a depth of coverage falling within the distribution of those containing BUSCO genes, as well as having one or more eukaryotic genes and/or transposable elements in the vicinity). Since the remaining 11.3% were scored either C (7.64%) or D (3.66%) (table S1), we considered candidates within the range A-D as valid candidates for endogenization. On the contrary, scaffolds annotated as F or X were rather considered as free-living viruses since they did not show eukaryotic genes or TE in their vicinity and had different coverage compared to BUSCO-containing scaffolds. Scaffolds classified as E were of unclear status and discarded.

#### Capacity to find domesticated EVEs (dEVEs)

An EVE was considered as domesticated if the dN/dS ratio was significantly below 1 or if TPM was above 1000. When dN/dS computations were possible (for 75/152 control EVEs), our pipeline considered the control EVEs as being under purifying selection in 70.39% of the cases. Overall, by combining the two metrics (dN/dS and TPM), our pipeline identified 69.04% of the control locus as being domesticated (table S1).

##### Capacity to infer events of endogenization (EVEs events)

Among the control species, the pipeline correctly inferred the expected 4 independent events: (1) *Leptopilina boulardi*/*Leptopilina clavipes*/*Leptopilina heterotoma* [19] (2) *Venturia canescens* [17], (3) *Fopius arisanus* [57], and (4) *Cotesia vestalis*/*Microplitis demolitor* [15] (table S1). However, in addition to the expected unique shared event concerning the *M. demolitor* and *C. vestalis* species, our pipeline inferred two additional events, each specific to one lineage. This was due to the fact that two genes were not detected by our pipeline as shared by *M. demolitor* and *C. vestalis*, either because they are effectively not shared (for 3 of them: HzNVorf118, like-*pif-4* (*19kda*), *fen-1*), or because of some false negative in one of the two lineage (for one of them:*p33* (*ac92*)).

#### Assessing the distribution of virus infecting insects

We estimated the number of viral species infecting insect species based on the virushostdb database (downloaded the 23/05/2022 on https://www.genome.jp/virushostdb/) which lists a wide diversity of viral species associated with their putative hosts. We also added two important exploratory RNA virus search studies [94, 100]. We kept only viruses found in interaction with insect in at least one of these databases. Genomic structures were retrieved through the ICTV report (2021.v1) and information available in ViralZone (all viral species details can be found on the GitHub repository under the name : All_virus_infecting_insects_informations.csv). We counted the number of viruses per genomic structure, and viruses from unknown genomic structures were discarded. In total, 2,626 viral species infecting insects were considered (detail : ssRNA(-) = 603sp, ssRNA(+) = 1,241sp, ssDNA = 75sp, dsRNA = 401sp, dsDNA =155sp, Unknown= 151sp). The Partiti-Picobirna, Narna-Levi, Mono-Chu, Bunya-Arenao, Luteo-Sobemo, Hepe-Virga and Picorna-Calici clades correspond to viral clades proposed by [94].

#### Divergence time estimation

We time-calibrated the inferred phylogenetic tree using a Bayesian approach on RevBayes 1.1.1 [101] and information on 5 fossils selected by [28]. Reduction of the supermatrix became necessary to overcome computational limitations when estimating node ages resulting from the large size of the concatenated BUSCO supermatrix (nsites = 228,009). We then generated one fasta file with a random draw without replacement of 20,000 sites from the supermatrix. Evaluation of the phylogenetic likelihood being the most expensive operation when calculating the posterior density, we decided to use the method developed in [102] to reduce computational cost and approximate the phylogenetic likelihood using a two-step approach. In the first step, the posterior distribution of branch lengths measured in expected number of substitutions is obtained for the fixed unrooted topology of using a standard MCMC analysis (100,000 iterations, 3 chains, 5000 burnin, tuningInterval=200). The obtained posterior distribution is then used to calculate the posterior mean and posterior variance of the branch length for each branch of the unrooted topology. In the second step, we date the phylogeny using a relaxed clock model and calibrations (500,000 iterations, 4 chains, 5000 burn-in, tuningInterval=200). Calibration of the root was done using a uniform law between 300 and 412 Mya. To verify that MCMC analyses converged to the same posterior distribution, for both steps we computed the effective sample size and applied the Kolmogorov-Smirnov test using the package convenience v1.0.0 with a minimum ESS threshold of 100 (however, due to an excessive demand for resources, we were unable to achieve the sampling value of an ESS of minimum 100 for 46/389 parameters (min=44.25)).

#### Ancestral state reconstruction

To explore the dynamics of EVEs gain in relation to lifestyle, we first had to reconstruct the ancestral lifestyle states of the Hymenoptera used in this study. This was achieved using a Bayesian approach implemented in RevBayes 1.1.1 [101]. The lifestyles of the Hymenoptera species used in this study were deduced from various sources (details and sources in the table named Assembly_genome_informations.csv from the GitHub repository). Since lifestyle characters are probably not equally likely to change from any one state to any other state, we choose the Mk model with relaxed settings allowing unequal transition rates. Thus, we assumed 6 different rates with an exponential prior distribution. Before running the MCMC chains, we made a preliminary MCMC simulation used to auto-tune the moves to improve mixing of the MCMC analysis with 1000 generations and a tuning interval of 300. We then ran two independent MCMC analyses, each set to run for 200 000 cycles, sampling every 200 cycles, and discarding the first 50 000 cycles as burn-in. To verify that MCMC analyses converged to the same posterior distribution, we computed the effective sample size and applied the KolmogoRov-Smirnov test using the package convenience v1.0.0 with a minimum ESS threshold of 100. The MCMC chain was subsampled to provide 1000 samples. At each sample, ancestral states were reconstructed for all nodes of the phylogeny. We assumed that the state assigned to a node was constant throughout the branch leading to that node.

#### Test of the lifestyle effect on viral endogenization and domestication

In order to test the lifestyle effect on the propensity to integrate and domesticate viral element, we first randomly sampled 1000 probable ancestral state scenarios to take into account the uncertainty in the estimates of the ancestral states of the nodes. Because a lot of branches had no EVE endogenization inferred, we ran zero-inflated negative-binomial GLM model, for each of these 1000 scenarios such that (GLM(NbEVEs∼free-living + endoparasitoid + ectoparasitoid * Branch_length, family = zero inflated neg binomial). We eliminated all branches older than 160 million years because they are too old for our analysis to detect events (the oldest event detected by our analysis is around 140 mya) that could artificially inflate the zero count. The model was implemented in stan language using the R package brms (seed = 12345, thin=5, nchains =4, niter = 10000) [103, 104]. Posterior predictive check was done using the package brmsfit in order to check that the model was correctly predicting the proportion of zeros. Indices relevant to describe and characterize the posterior distributions were computed using the R package BayestestR [105]. Autocorrelation was studied using the effective sample size index (ESS) with a value greater than 1000 being sufficient for stable estimates [103]. The convergence of Markov chains was evaluated by a Rhat statistic equal to 1. All the posterior coefficient estimated values were then pooled together (after checking the convergence of all chains via the GelmanRubin function in R [106]) and compared between the free-living, endoparasitoid and ectoparasitoid modalities.

To calculate the rate of domestication independent of the rate of endogenization, we built a binomial logistic regression model in a Bayesian framework, specifying the number of domesticated EVEs (or Events) (the numerator) relative to the total number of EVEs or Events inferred by our pipeline (the denominator). These binomial models allowed us to test whether the probability of domestication after endogenization correlated with lifestyle by controlling for the endogenization input (the denominator). Thus, for each of the 1000 lifestyle scenarios, we ran a binomial brms model with a logit link such that brms(Nb dEVEs/dEvents | trials(Nb EVEs/Events) ∼lifestyle + Branch length).

Before analyzing the data, we checked that the inferences did not depend on the quality of the genomes selected for analysis. We found a significant effect of the lifecycle on the N50 and percentage of complete+partial BUSCO in the assemblies (Kruskal-Wallis rank sum test p-values respectively = 3.192e-10 and 1.26e-14). Furthermore, a pairwise Wilcox test with p-values adjusted with Bonferroni method revealed a significantly higher values of N50 and %complete+partial BUSCO in genome assemblies from free-living species compared to endo and ectoparasitoids species (p-value <0.05). The same test using the total assembly length in bp did not reveal any difference between the three lifestyles (p-value >0.05). Overall, free-living species have better assemblies. Because better assembly quality should facilitate the discovery of endogenous viral elements both by sequence homology detection and by a better assessment of the endogenized nature of the EVE (scaffolds A,B,C and D), we should thus underestimate the number of EVEs in endo- and ectoparasitoid species compared to free-living species. Since our analysis led to opposite conclusion, our results cannot be explained by this feature of the dataset.

## Author contributions

BG & JV conceptualized the study. BG coordinated and performed all bio-informatic analysis, wrote the first draft, and edited the manuscript. BB participated in the Bayesian analysis. DDV developed the scripts used in the phylogenetic analysis to infer endogenization events. SC, DGN, R. J-Y, S.J, S.M, R.N, C.S, N-B.P and J-T.E provided the specimens. DL did all the molecular biology work. BG, SC and JV participated in the interpretation of the results. BG, SC & JV wrote the manuscript and integrated the comments done by all co-authors.

## Data availability

All sequencing data are available at NCBI via the BioProject accession number NCBI: PRJNA826991. All scripts and additional tables as well as all cluster phylogeny figures are available under the github repository : (BenjaminGuinet/Viral-domestication-among-Hymenoptera).

## Acknowledgments

This work was performed using the computing facilities of the CC LBBE/PRABI. We thank Clément Gilbert and Jean-Michel Drezen for helpful discussions and Elijah J Talamas, Nicolas Ris, Lene Sigsgaard for providing specimens. This work was supported by a grant from the Agence Nationale de la Recherche (ANR) to J.V. (11-JSV7-0011 Viromics). We also thank the ANR project HORIZON (SC) for financial support.

## Declaration of interests

The authors declare no competing interests.

## Supporting Information

### 1- ssRNA endogenization

Although our results show an under-representation of EVEs deriving from ssRNA viruses (relatively to their high abundance in insect virome), they were involved in a high absolute number of endogenization events: 21 viral families/clades were involved in 174 independent endogenization events in the 114 Hymenoptera genomes analyzed, in particular involving *Chuviridae*, *Artoviridae* and *Nyamiviridae* (Figure2-B). In a recent meta-analysis [4], more than 1876 EVEs involving ssRNA viruses were identified in 37 species distributed in 8 insect orders. Interestingly, the authors noticed that the contribution of negative-stranded RNA viruses was overall high (67%), but was also highly species-specific. In our dataset, the great majority of ssRNA viruses donors were negative-stranded (14/21 viral family/clades) accounting for 78.8% of ssRNA events. The pattern thus seems even more pronounced in Hymenoptera compared to insects in general, and resemble the pattern observed in ticks [56]. The reasons for the asymmetry observed between negative and positive strand RNA viruses in endogenization are unclear. One explanation proposed by [107] posits that the ssRNA(-) have a higher probability of endogenization compared to ssRNA(+) because non-segmented ssRNA(-) usually produce abundant short mRNAs compared to ssRNA(+) which conversely produce lower amount of long mRNAs encoding a single polyprotein [108]. Then, all else being equal, an RNA(-) virus would produce more RNA molecules, which will increase the likelihood that some of them get reverse-transcribed and ultimately endogenized into the host genome. In support of that view, [107] noticed that the NP gene was more often endogenized compared to the other genes encoded by most ssRNA(-), which fits its prediction. This is because for most Mononegavirales species the 3’ nucleoprotein (NP) gene is the most abundant RNA [109], due to the polar 3’-5’ stepwise attenuation of transcription [109]. This pattern was since observed on some systems (i.e. mosquitoes and few mammals genomes [1]) but opposite results were also obtained on others (ticks, [56]). In our dataset, we found two ssRNA(-) non-segmented families showing the expected pattern where the genes closest to the 3’ regions were the most endogenized : the Nucleoprotein (N) which is the first transcribed protein in *Nyamiviridae* was endogenized in 26 cases out of 28, and the 3’-unknown protein in *Lispiviridae* (first transcribed) was endogenized in 12 cases out of 15 endogenization events (FigureS3). On the contrary, all the other putative ssRNA families donor presented more EVEs deriving from the middle or the 5’ genomic regions: the most endogenized gene from *Artoviridae* was the U2 protein (19/39) which is in the middle of the genome (2nd/3); in *Bornaviridae* and *Rhabdoviridae* the most endogenized gene was the RdRp (L) protein, which is the last ORF in the first genome (out of 8 genes) and the one just before the last gene in *Rhabdoviridae*. Finally, out of 36 EVEs deriving from *Chuviridae*, 26 corresponded to the Nucleoprotein (N) which is the last transcribed protein in the closest viral genomes. These unexpected results under Holmes model may thus lead one to reject the hypothesis, unless peculiar mechanisms of regulation of the transcription are at play for these viruses. Another explanation could come from a strong selective pressure for retaining particular proteins (i.e. Nucleoprotein) in the genome, independently of their level of transcription.

### 2- A new case of virus domestication in Platygaster orseoliae

In the assembly of P. orseoliae, 12 scaffolds were annotated as free-living viruses (F-X scaffolds). They had a different sequencing depth compared to BUSCO containing scaffolds and encoded 136 complete ORFs for which 21 presented homology with LbFV ORFs (min bit score = 50, min ORF size = 150pb, max overlaps = 23pb). ORF density was 82.7% which is in the range of expected values for related free-living viruses [110]. In order to identify additional scaffolds possibly belonging to this free-living virus, we searched for homology between the 136 putative viral proteins, and the scaffolds obtained from the assembly of *P. orseoliae*. These results allowed us to identify two additional scaffolds (scaffold_117128 scaffold_18896). Because the total size of the 14 putative “free-living” scaffolds was within the expected range for a dsDNA virus genome (136,801 bp) and because the average coverage was much higher than BUSCO-containing scaffolds (mean cov = 95.6X [sd=5.05X] compared to 33X in BUSCOs) and homogeneous (Figure S4), we believe that this set of scaffolds corresponds to the whole genome of a new virus, related to LbFV, which we propose to call Platygaster orseoliae filamentous virus (PoFV). In order to characterize the relationship of this new virus within dsDNA viruses diversity, we inferred a phylogenetic tree including ORFs of known dsDNA viruses along with the EVEs newly identified here. The phylogenetic reconstruction revealed that PoFV was the closest relative of the endogenized virus found in the same species (PoEFV, Figure 5).

In order to detect possible new viral endogenization from the same “donor virus”, we queried the genome of *P. orseoliae* with the 136 predicted proteins of PoFV. This way, we found a total of 139 convincing hits (89 PoFV ORFs), including the hits to the 15 ORFs with LbFV-homology. All ORFs were encoded by scaffolds with BUSCO-like coverage depth (p-value cov >= 0.05) and/or containing eukaryotic genes and/or transposable elements (Blastx E-value max = 7.060e-07, bits min =50, with an average percentage of identity of 69.16%). Furthermore, a large proportion of the EVEs (22.7%) presented premature stop codons within the sequences, further suggesting that these virally-derived genes are indeed endogenized since abundant pseudogenization is not expected in free-living virus genomes (Figure S8-A).

Among the 81/139 apparently intact EVEs (with ORF length >= 50% of the PoFV ORF), some are likely implicated in DNA replication (*integrase*), in transcription (*lef-8*,*lef-9*,*lef-5*,*lef-4*), in packaging and envelopment (*ac81*, *38k*) and in infectivity (*pif-1*, *pif-2*, *pif-3*). Among the 139 PoFV-related EVEs found in *P. orseoliae*, 104 corresponded to putative paralogs. Conversely, none of these 104 ORFs were present in two copies within the PoFV genome, suggesting that a major post-endogenization duplication event occurred or that multiple endogenization events did occur. Among these 104 duplicated EVEs, 78 presented topologies allowing us to calculate dN/dS ratios using Bayesian pairwise estimates (runmode -3 in codeml) or foreground/background tests (codeml) when topologies presented more than 4 leafs. Before running the foreground/background tests, we constrained all paralogs to form a monophyletic group including the PoFV loci as the closest taxa in the phylogenies (all LRT tests did not significantly present differences between constrained and unconstrained trees). Among these 78 paralogs EVEs, 44 presented a complete and intact open reading frame and a dN/dS ratio significantly lower than 1 suggesting that they are under stabilizing selection (Figure S8-A).

Although functional studies are needed to confirm that these virus-derived genes are involved in the formation of VLPs as observed in *Leptopilina* [19], we believe that *P. orseoliae* filamentous virus (PoEFV) is a good candidate for viral domestication, possibly involved in counteracting its Diptera host immune system (from the family Cecidomyiidae [33]).

### 4- Assignation of the unknown Hymenoptera to species

UCEs along with 400 bp of flanking regions on either side were extracted from the low coverage scaffolds with a custom script. We used a two-step process to assign the unknown sample to species. First, UCEs + flanking regions were analyzed with a set of UCEs + flanking regions obtained from early diverging families of Chalcidoidea by [70, 69] to assign the unknown sample to family. Then, unknown sequences were analysed with a larger set of species belonging to the identified family (Eulophidae; loci taken from [69]). In both cases, only loci that had a sequence for at least 75% of the samples included in the analysis were retained. Loci were aligned with MAFFT (-linsi option; [111]). Positions with > 90% gaps and sequences with > 25% gaps were removed from the alignments using SEQTOOLS (package PASTA; [112]). The concatenation of all loci was analysed with IQ-TREE v 2.0.6 [71] without partitioning. Best fit models were selected with the Bayesian Information Criterion (BIC) as implemented in ModelFinder ([72]). FreeRate models with up to ten categories of rates were included in tests. The candidate tree set for all tree searches was composed of 98 parsimony trees + 1 BIONJ tree and only the 20 best initial trees were retained for NNI search. Statistical support of nodes was assessed with ultrafast bootstrap (UFBoot) ([113]) with a minimum correlation coefficient set to 0.99 and 1,000 replicates of SH-aLRT tests ([114]). Results of the phylogenetic analyses are presented in Figure S2. Placement of the unknown species in trees shows that samples of *P. orseoliae* were likely mixed up with a species belonging to the genus *Aprostocetus* (Eulophidae, Tetrastichinae). Given its small size, color and abundance (265 species described just in Europe), it seems plausible that one specimen of *Aprostocetus* sp. remained unnoticed in the pool of *P. orseoliae*.

---

**Figure S1.**
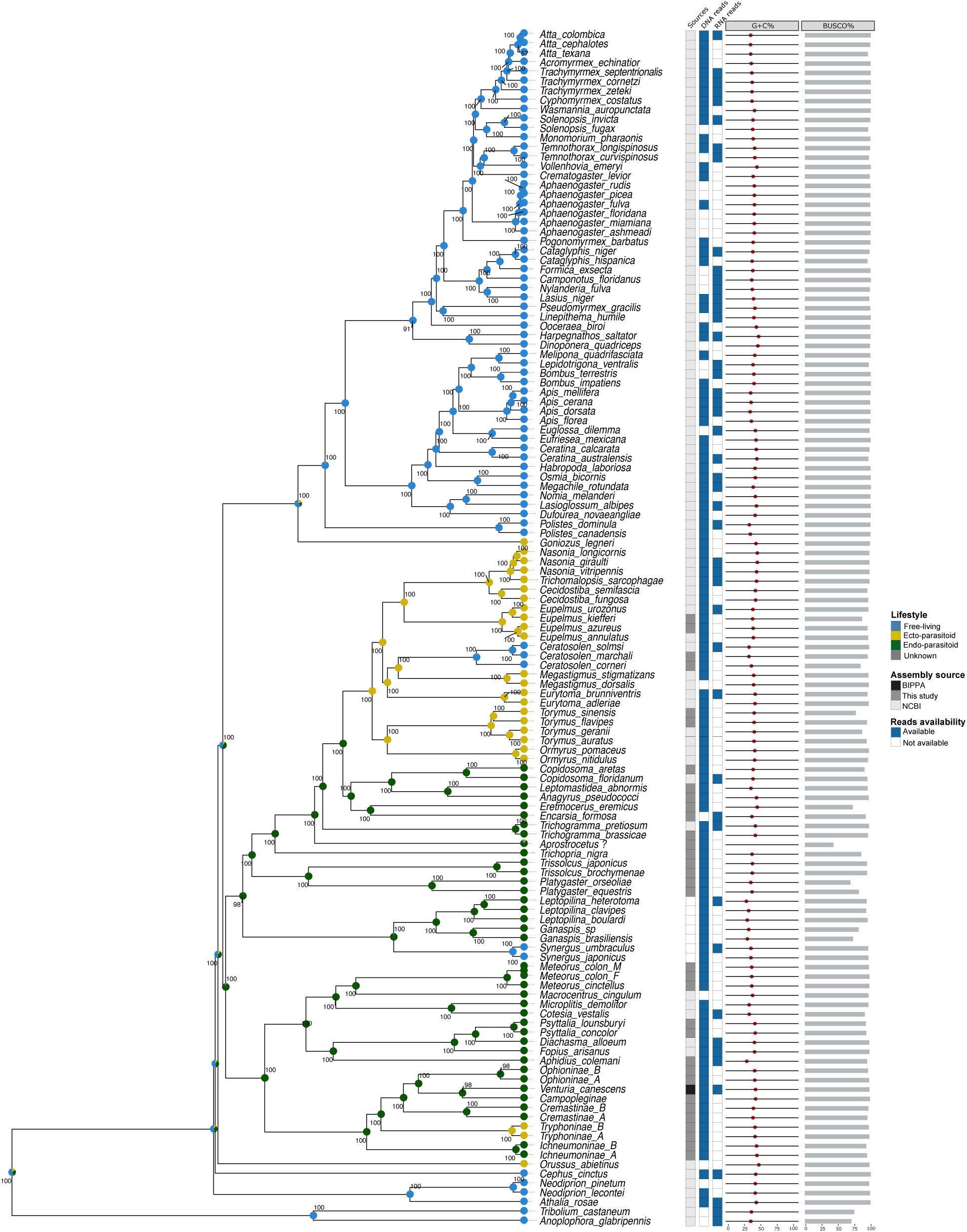
Source of the datasets and availability of the reads. Phylogeny of 124 Hymenoptera species. Two Coleoptera species were used to root the tree. The aLRT bootstrap scores are represented along the nodes. The sources refer to the platform or laboratory in which the assemblages come from (This study, BIPPA: BioInformatics Platform for Agroecosystem Arthropods, NCBI: National Center for Biotechnology Information). The assemblies for which raw DNAseq or RNAseqs reads were available are listed in the column DNA or RNA reads. The G+C% column reflects the average G+C rate for each assembly, and the BUSCO% column reflects the rate of complete or partial BUSCOs found via the analysis with BUSCO V3. Posterior Bayesian lifestyle inference distribution for each node and tips are represented by colored pie charts.

**Figure S2.**
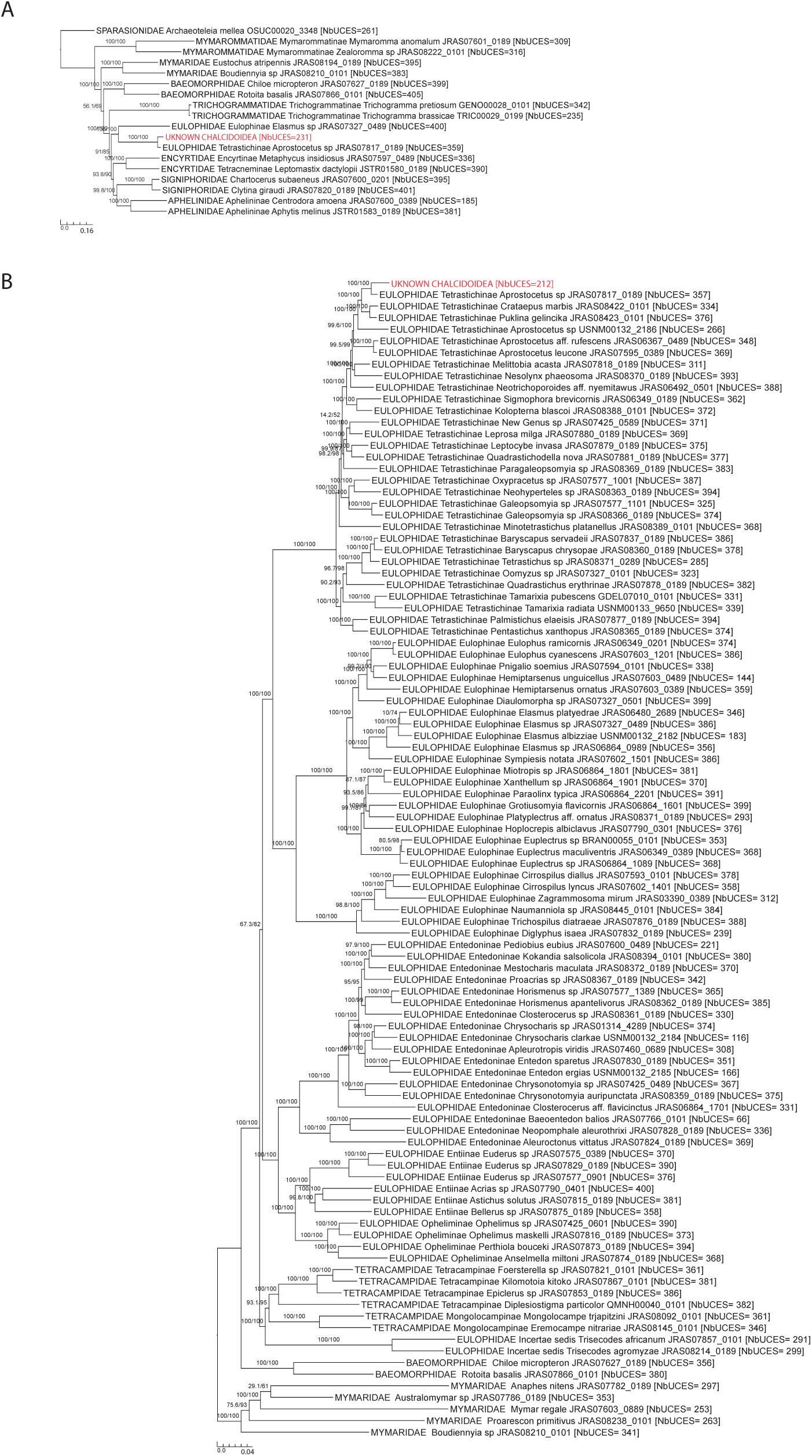
UCE trees built to assign to species the unknown Chalcidoidea sequenced with the pool of *P. orseoliae*. **A**: Phylogeny of early diverging families of Chalcidoidea (423 UCES and 127,979 bp were analysed to get the tree, best fit model = GTR+F+R10). **B** : Phylogeny of the family Eulophidae to which the unknown sample was inferred to belong to (408 UCES and 77,514 bp were analysed to get the tree, best fit model = GTR+F+R10). For both trees, SH-aLRT/UFBoot are shown at nodes; the number of UCEs analyzed for each sample is indicated between bracket and the unknown sample is highlighted in red.

**Figure S3.**
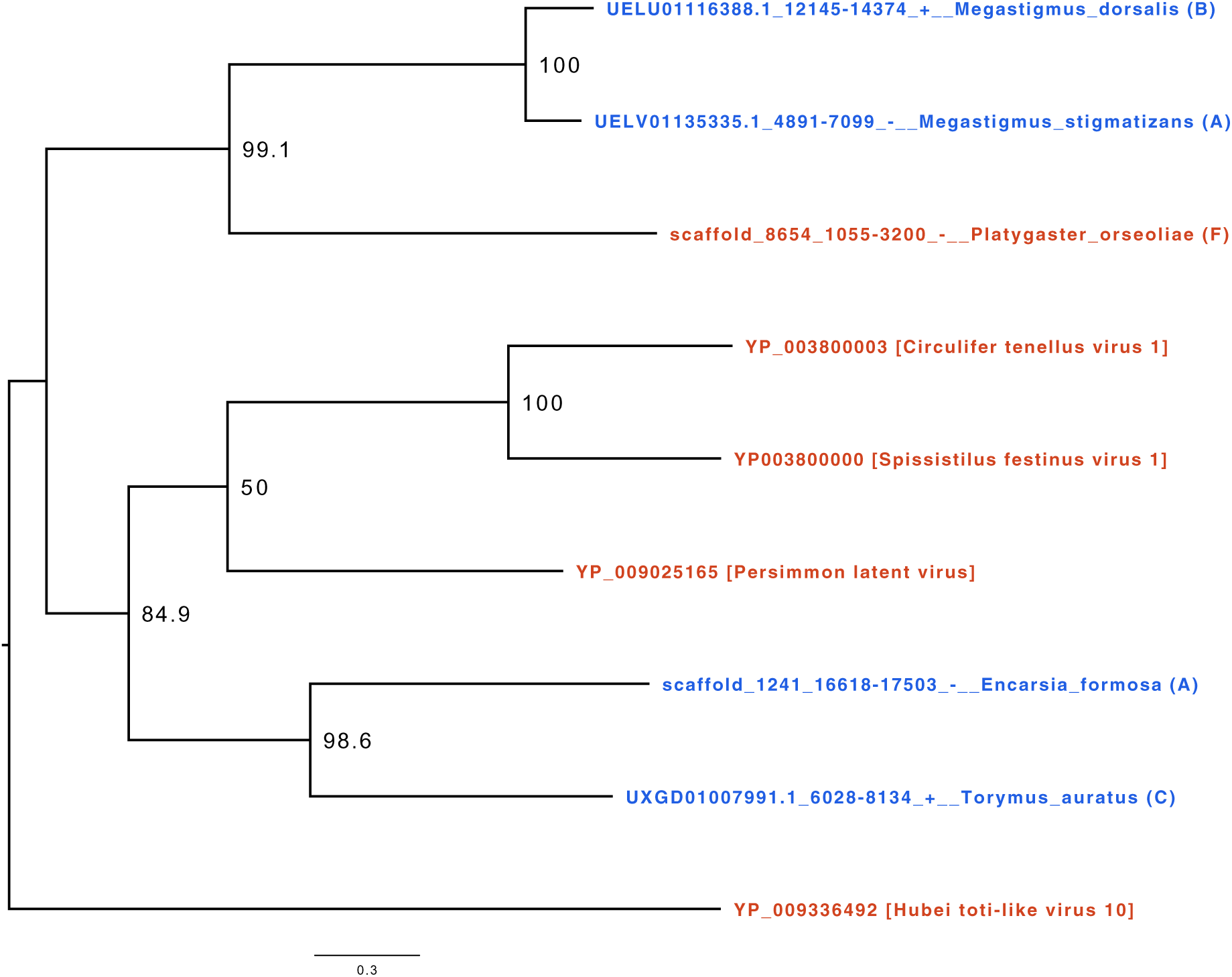
Example of endogenization events. The phylogeny of cluster21304 corresponds to the clustering of a set of viral and candidate viral insertion genes sharing a homology. In red are represented the loci of viral origin, and in blue are represented the loci probably endogenized (EVEs). The letter at the end of the taxon label represents the endogenization score assigned to the candidate (see details in Materials and methods). In this example, we found two singular endogenization events in the species endoparasitoid *Encarsia formosa* (annotated A and thus presenting a depth of coverage non-significantly different from the distribution of the BUSOs of the genome as well as at least one transposable element and/or one eukaryotic gene) and ectoparasitoid *Torymus auratus* (annotated C and thus presenting only a depth of coverage non-significantly different from the distribution of the BUSOs of the genome). Since these two species do not share a close common ancestor in the phylogeny and come from two different families, the algorithm therefore assigned them to two independent viral endogenization events. The viral locus found in the assembly of the endoparasitoid species *Platygaster orseoliae* was annotated F, meaning that the depth of coverage deviated significantly from the BUSCO distribution of the genome and that no TEs and less than 5 eukaryotic genes were found in the scaffold containing the candidate insertion. Finally, the two loci belonging to the ectoparasitoid species *Megastigmus stigmantizans* and *Megastigmus dorsalis* both show a score supporting viral endogenization. Furthermore, these species exhibit a doubly monophyletic clade (high bootstrap score) within the gene phylogeny and within the species phylogeny, suggesting that they acquired this viral gene from their closest common ancestor about 20 million years ago. All newick phylogenies are available on the GitHub repository under the name : All*_c_ luster_p_hylogenies_m_erged*.

**Figure S4.**
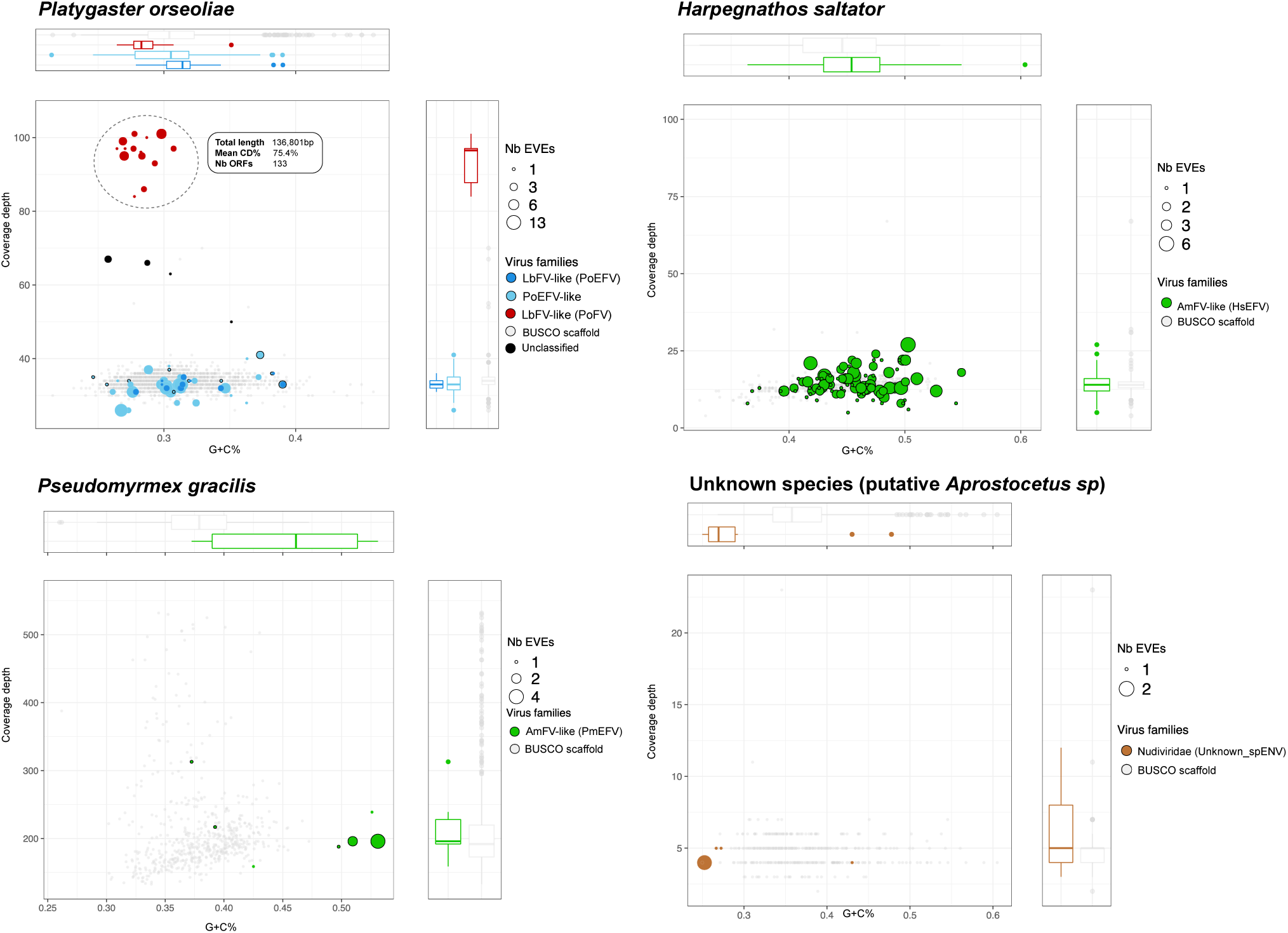
G+C%Coverage distribution of scaffold containing multiple EVEs Events. The size of the dots corresponds to the number of candidate EVEs inside the scaffold. The color represents the genomic entity from which the EVE probably originated (brown: *Nudiviridae*, blue = LbFV and green = AmFv). The red color refers to scaffolds showing free-living virus signatures. The grey color refers to scaffolds containing one or more BUSCO genes. The dots circled in black correspond to scaffolds that contain one or more eukaryotic genes and/or repeat elements.

**Figure S5.**
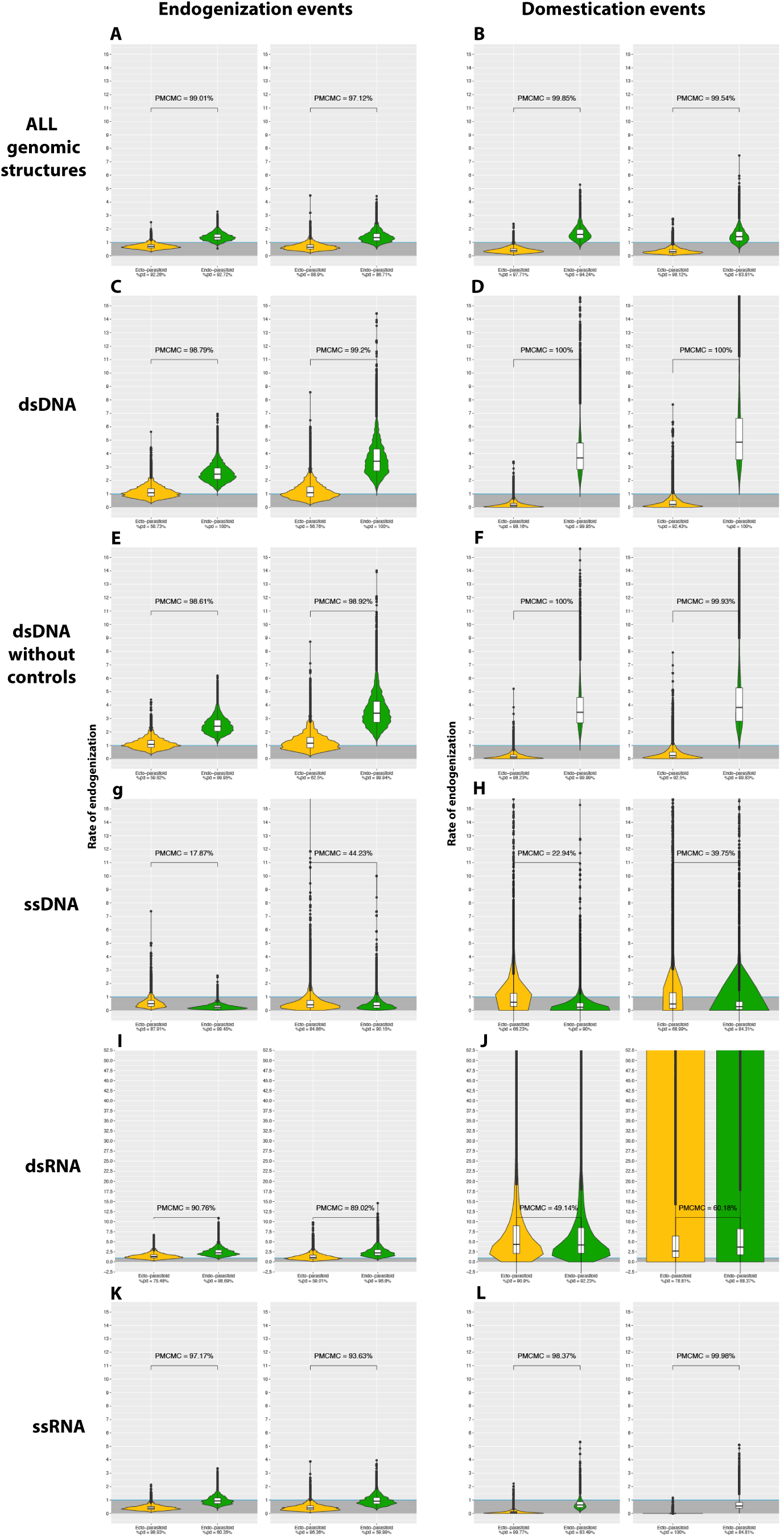
Violin plots of the posterior distribution of GLM coeficients after exponential transformation in relation to wasp lifestyle. The ectoparasitoid lifestyle is in yellow, the endoparasitoid lifestyle is in green, and the free-living lifestyle is in blue. Coefficients have been transformed into exponential and correspond to the posterior distribution of the coefficients of a binomial negative zero-inflated GLM model, where the lifestyle free-living stand for the intercept. The Y-axis corresponds to the multiplicative factor of the number of endogenization and/or domestication of EVES and/or events relative to free-living species. The coefficients are derived from 1000 GLM models adjusted on 1000 randomly selected probable scenarios (>90 CI) of ancestral states at nodes. Branches from nodes older than 160 million years have been removed from the dataset. The ROPE% is the percentage of the posterior distribution of coefficients below the intercept. The posterior distribution of the interaction coefficients between lifestyles and branch size were

**Figure S6.**
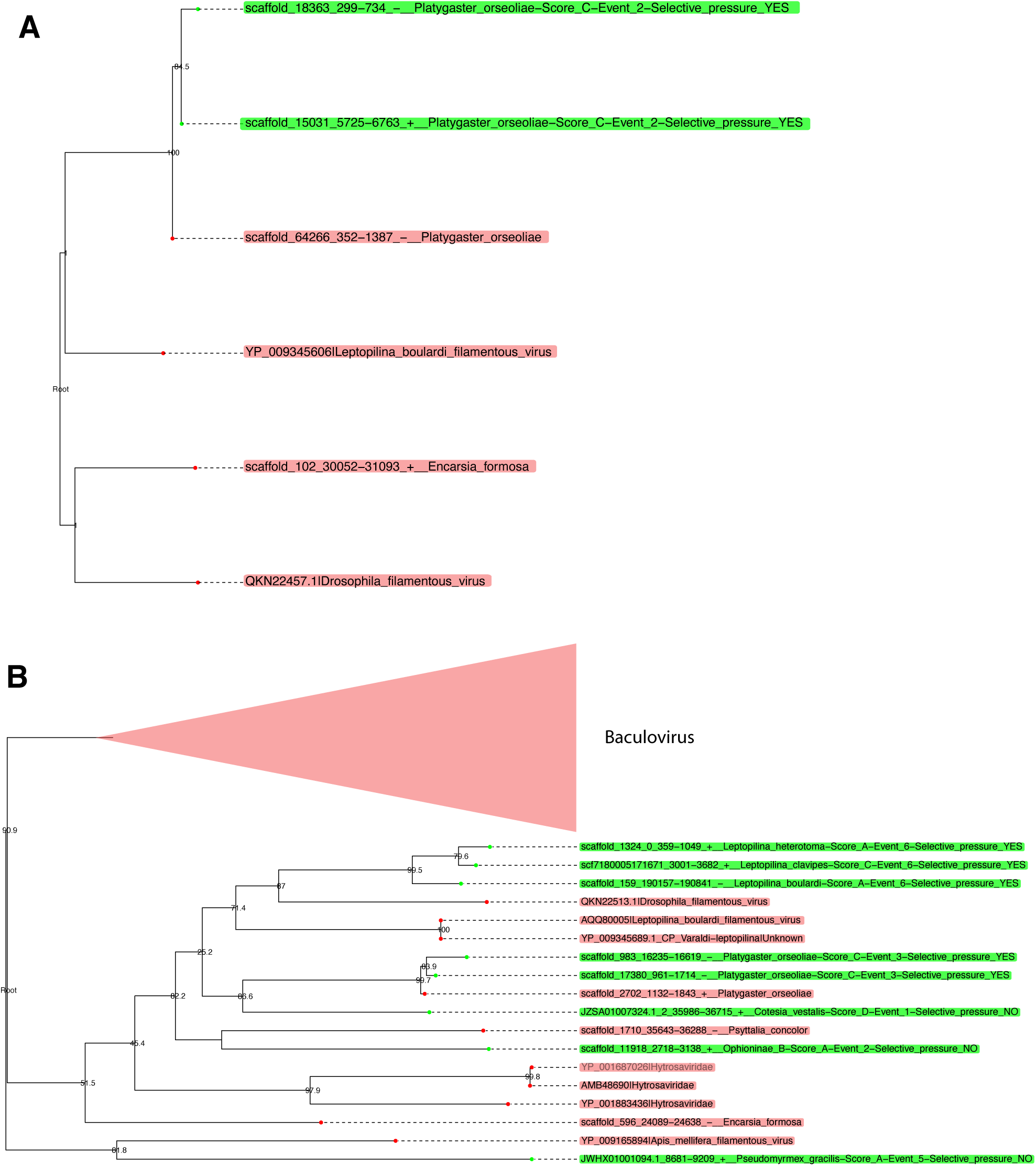
Phylogenies of LbFV-like proteins under purifying selection in *Platygaster orseoliae* genome. The panel A represents the Cluster_25710 which corresponds to the *integrase* protein. The panel B represents the Cluster_26675 which corresponds to the *ac81* protein. Taxa in red correspond to putative loci belonging to free-living viruses, while green taxa correspond to putative EVEs (the assigned family is indicated after the pipe). Green labels indicate the following information : (1) scaffold name, (2) start location, (3) end location, (4) strand, (5) Hymenoptera species, (6) endogenization index, (7) the inferred event number within the cluster phylogeny, and (8) whether the loci are found under purifying selection (YES) or not (NO).

**Figure S7.**
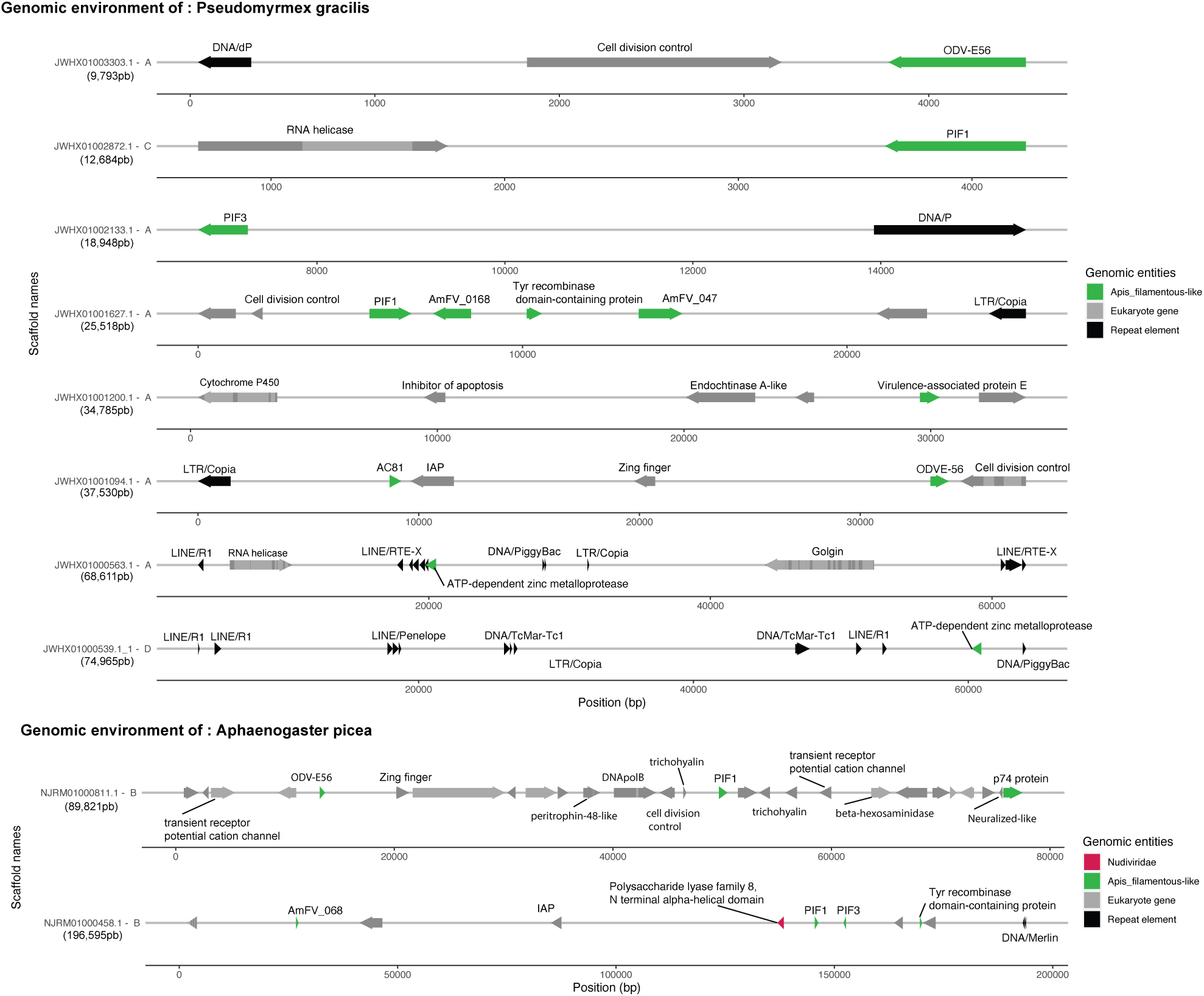
Candidate EVEs in two ant species. In grey are displayed the eukaryotic genes predicted by Augustus, with a dark color for exons, and light for introns. In black are displayed the transposable elements predicted by sequence homologies with the RepeatPeps protein database. The size of the scaffold is displayed below the name of each scaffold. The letter followed by the scaffold name refers to the scoring given to the scaffold based on coverage and gene/ET presence information (see details in Materials and methods). For the sake of representation, all scaffolds are represented at the different scale, the exact coordinates of the elements are referred in the abscissa which corresponds to the coordinates in base pairs. Annotation is indicated above the arrows.

**Figure S8.**
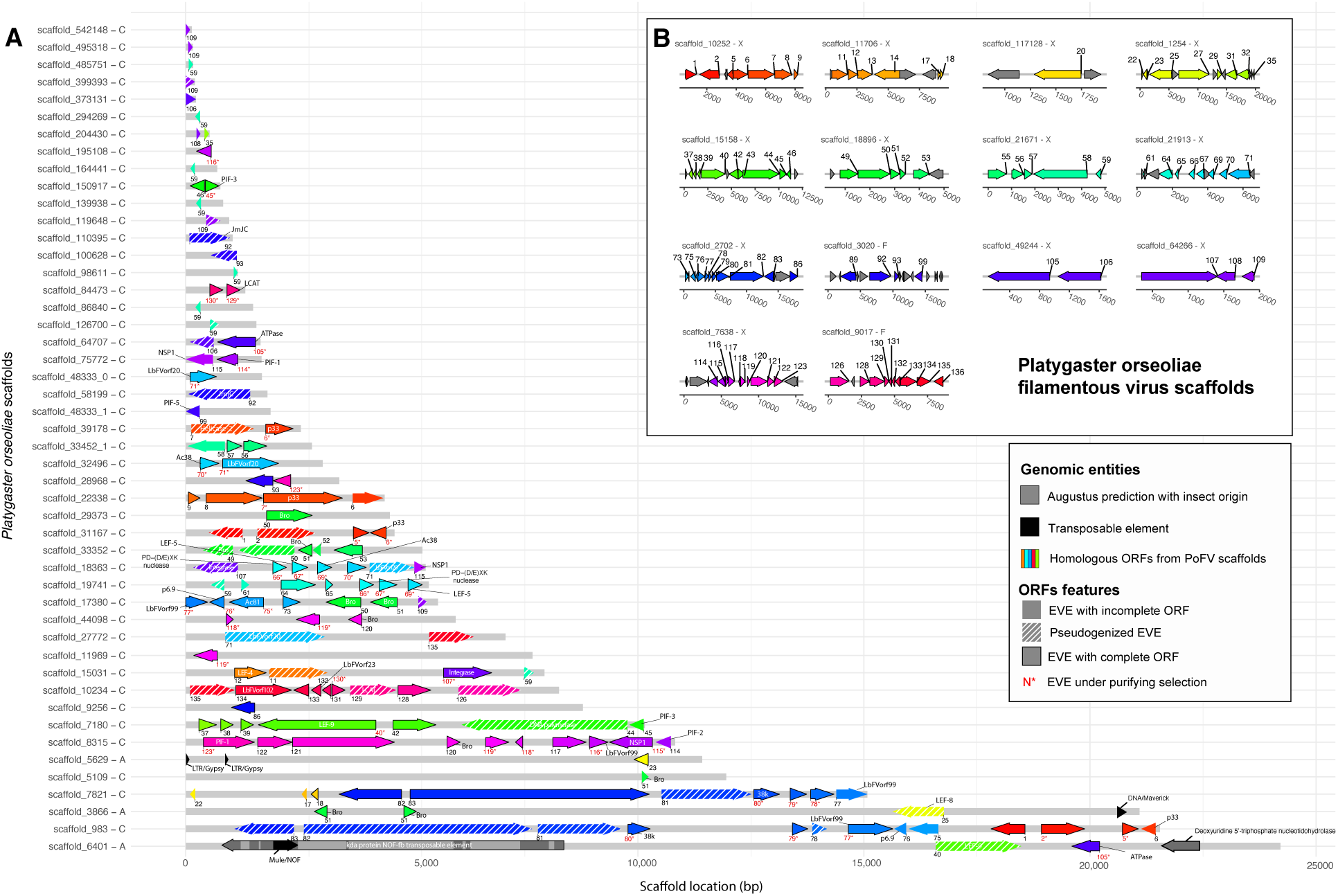
Genomic environment for the EVEs detected in *Platygaster orseoliae*. The plot show regions homologous to viral ORFs in the *Platygaster orseoliae ilamentous virus* (PoFV) genome (A). The colored regions correspond to the predicted ORFs in the PoFV genome (gray ORFS in PoFV scaffolds mean that no homologous EVEs were found in the genome of *P.orseoliae*.) (B), so the closer the colours, the closer the ORFs were initially in the PoFV genome. In grey are displayed the eukaryotic genes predicted by Augustus, with a dark color for exons, and light for introns. In black are displayed the transposable elements predicted by sequence homologies with the RepeatPeps protein database (max E-value =5.866e-18). The letter followed by the scaffold name refers to the scoring given to the scaffold based on coverage and gene/ET presence information (see details in Materials and methods). The exact coordinates of the elements are referred to the x abscissa, which corresponds to the coordinates in base pairs and can be found on the Github repository under the name : All_PoFV_PoEFV_loci_informations.txt. Annotation is indicated in or next to the arrows. The number below each EVE correspond to the homologous ORF number in the PoFV genome. Numbers are colored in red if the EVE has at least one paralog and if the computed dN/dS is below 1, suggesting purifying selection in the *P.orseoliae* genome. Arrow with black borders correspond to EVEs showing a complete ORF (>50% of the size of the best PoFV ORF). Hatched arrows correspond to pseudogenized EVEs (with premature stop codons). The colour difference between black and white for the names of the proteins is for visual purposes only.

**Figure S9.**
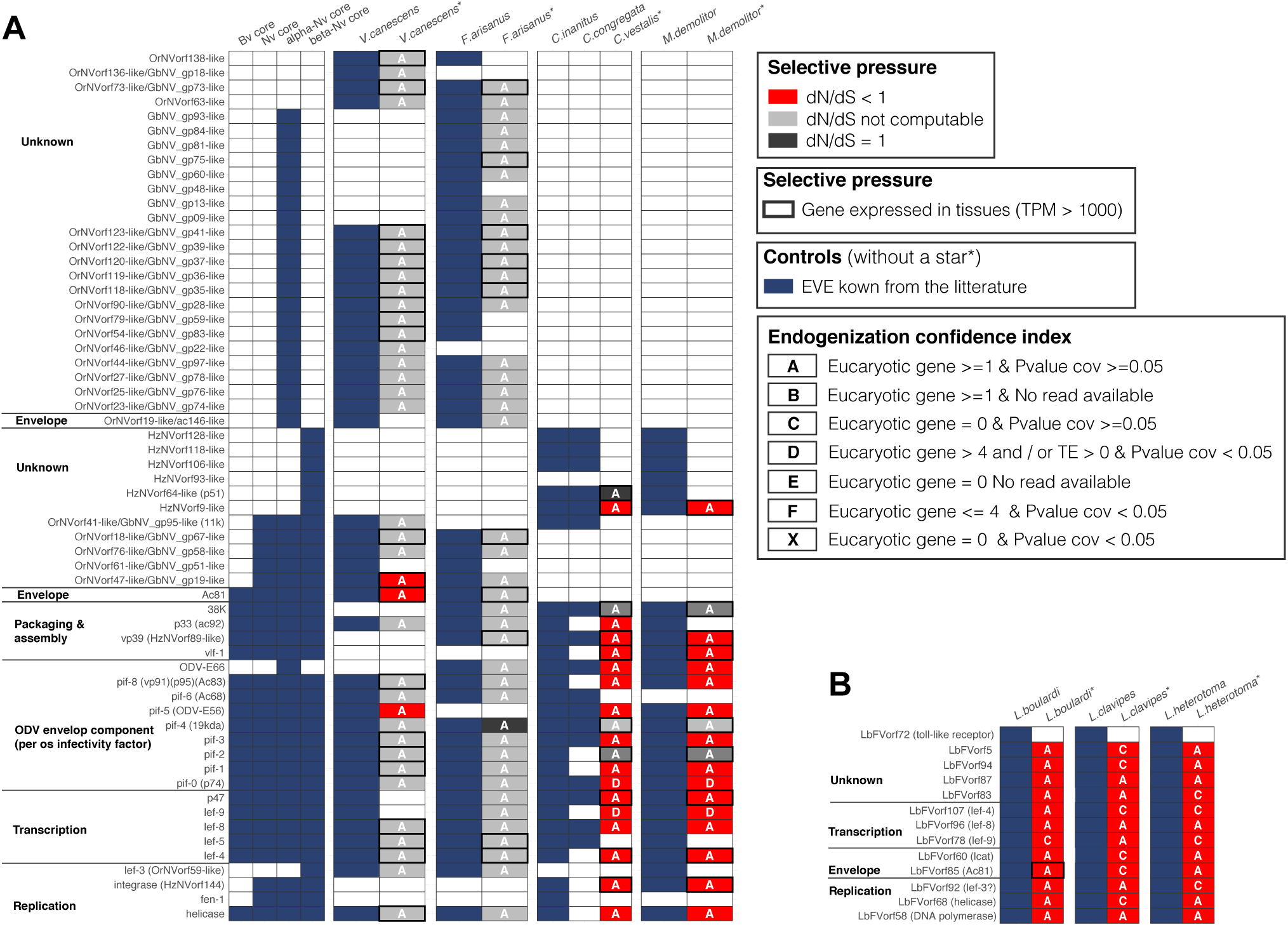
Heatmap representing the viral genes known to be domesticated by Hymenoptera. The panel (A) refers to the four known cases (*Venturia canescens*, *Fopius arisanus*, *Cotesia congregata* and *Microplitis demolitor*) involving Nudivirus donors while the panel (B) refers to the known case involving LbFV donors in three *Leptopilina* species. Complete parasitoid wasp genomes information were available for *Microplitis demolitor*, *Venturia canescens*, *Fopius arisanus* and *Cotesia congregata*, while only partial genomic data were available for *Chelonus inanitus*. Each row indicates a gene which has been identified previously as being endogenized in at least one species. In (A), the first four columns indicate whether the gene is a core gene for baculoviruses (Bv), Nudiviruses (Nd), alpha-nudiviruses (alpha-Nv) or beta-nudivirus (beta-nv). The following columns indicate the presence of each gene based on the literature (in blue) and based on our pipeline (columns with a star symbol). The colors indicate the inferred selection pressure acting on each gene (dN/dS) and the letters A, B, C, D, E, and X represent the degree of confidence in the endogenization. Capital letters indicate that this gene is present in a scaffold that contains other candidate genes. When the box is framed in black, it means that the gene is expressed (TPM>1000).

**Figure S10.**
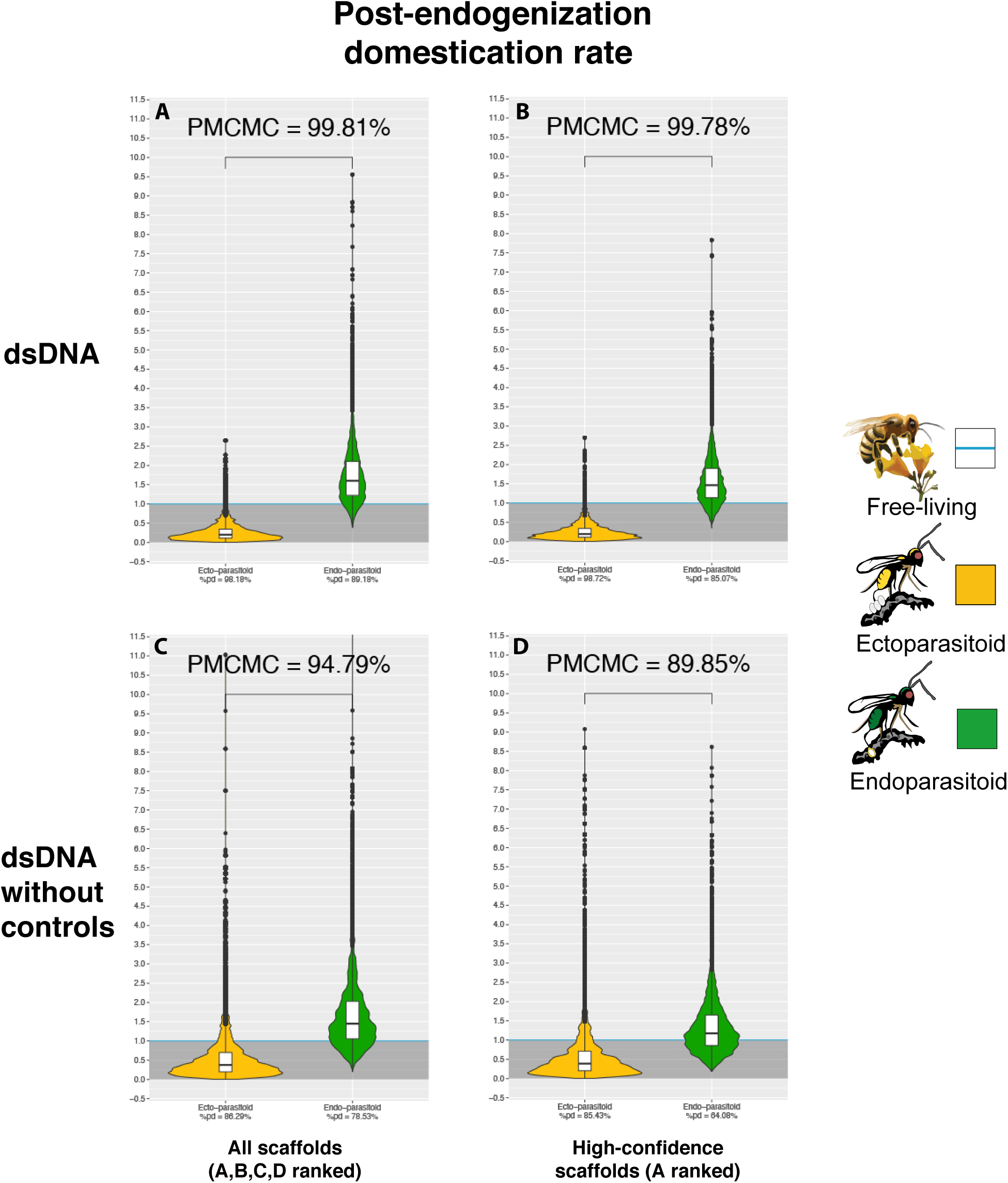
Violin plots of the posterior distribution of dEvents GLM coeficients in relation to wasp lifestyle (corrected for Events rates). The ectoparasitoid lifestyle is in yellow, the endoparasitoid lifestyle is in green, and the free-living lifestyle is in blue (the intercept). Coefficients have been transformed into exponential and correspond to the posterior distribution of the coefficients of a binomial logistic regression GLM model, where the lifestyle free-living stand for the intercept. The Y-axis corresponds to the multiplicative factor of the number of dEvents (corrected for Events rates) correlative to free-living species. The coefficients are derived from 1000 GLM models adjusted on 1000 randomly selected probable scenarios (>90 CI) of ancestral states at nodes. Branches from nodes older than 160 million years have been removed from the dataset. The ROPE% is the percentage of the posterior distribution of coefficients below the intercept. The posterior distribution of the interaction coefficients between lifestyles and branch size were not informative, and the branch size factor was therefore added as an additive effect to the model. **A**- The corrected coefficient within all dEvents, **B**- The corrected coefficient within all dEvents without the control genomes, **C**- The corrected coefficient within all dEvents present in a scaffold annotated with a score A, **D**- The corrected coefficient within all dEvents present in a scaffold annotated with a score A and without the control genomes.

**Figure S11.**
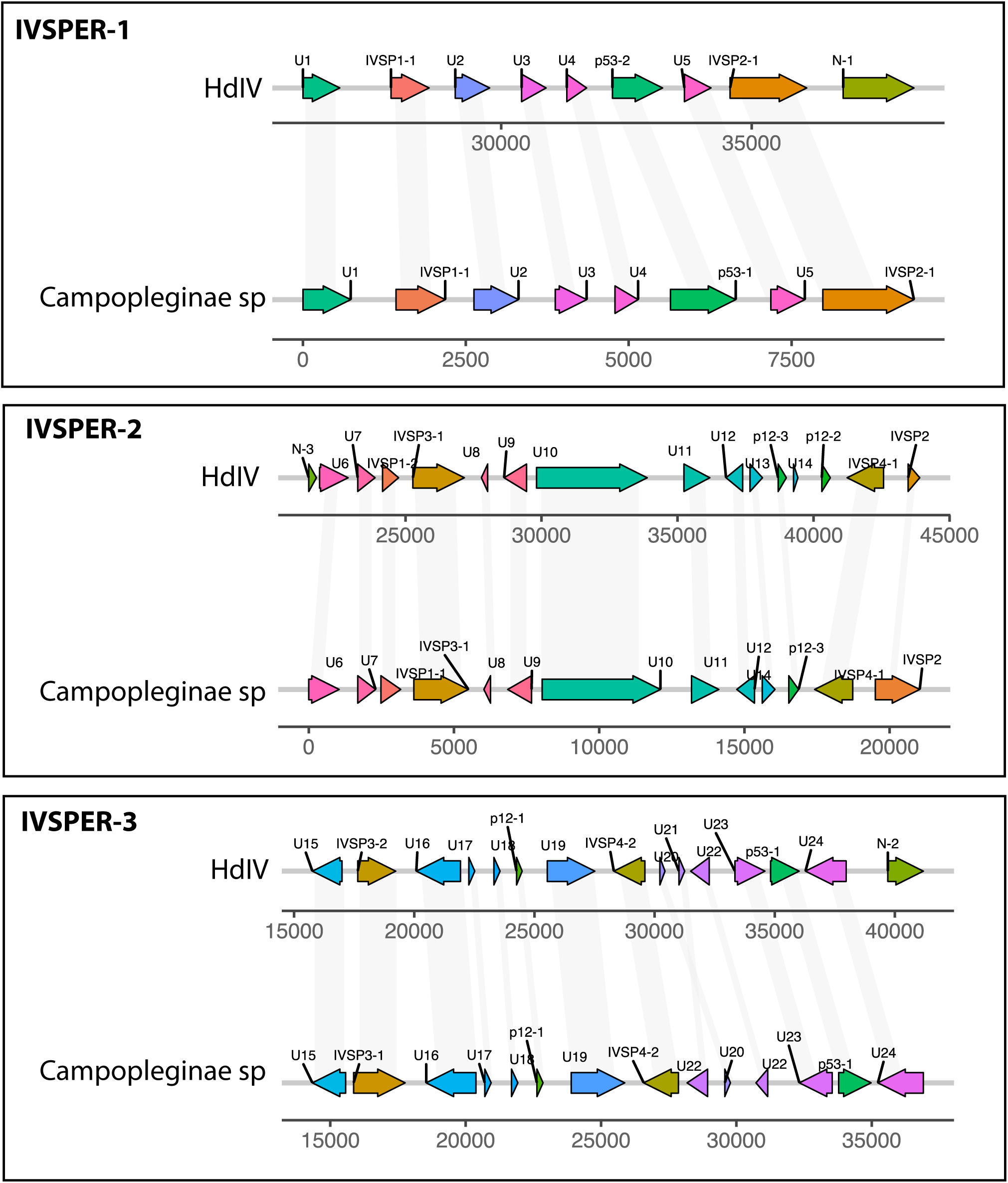
IVSPER genes identified in the Campopleginae genome. The figure compares the synteny of the IVSPER between Hyposoter didymator ichnovirus (HdIV) and the Campoplegninae of our dataset. Homologous genes with synteny between the two species are indicated by grey shading. The direction of the arrows corresponds to the sense and anti-sense strand. The color of the boxes is unique to each IVSPER.

**Figure S12.**
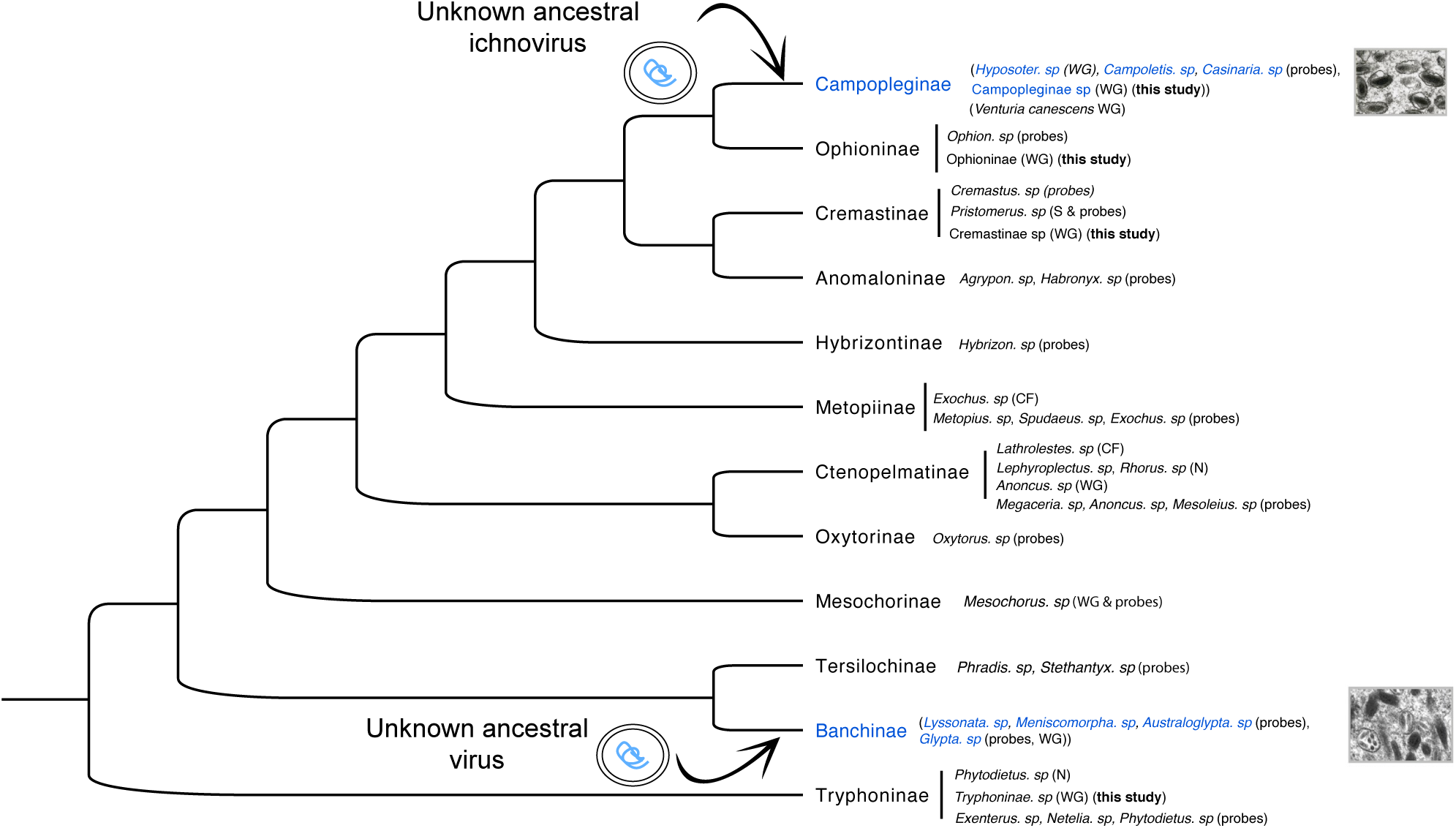
Cladogram of the Ophioniformes group, illustrating the two independent endogenization events of two unknown viruses in Banchinae and Campopleginae lineages. The phylogeny includes 12 subfamilies of the Ophioniformes group within the superfamily Ichneumonoidea. Several species of these subfamilies have been examined for the presence of ichnovirus-like polydnaviruses: by negative staining of calyx fluid (N), TEM of ovarian sections (S), visual examination of the calyx fluid (CF), probes from ichnovirus replication or structural proteins (probes) or by IVSPER sequence homology on whole genome assemblies (WG). The subfamilies and species in blue correspond to those positive for a dsDNA virus endogenization from unknown origin (ichnovirus-like). The others (in black) were negative for endogenized ichnovirus-like elements. The phylogeny is inspired from [115], and the data reported comes from [115, 116, 117, 118].

**Figure S13.**
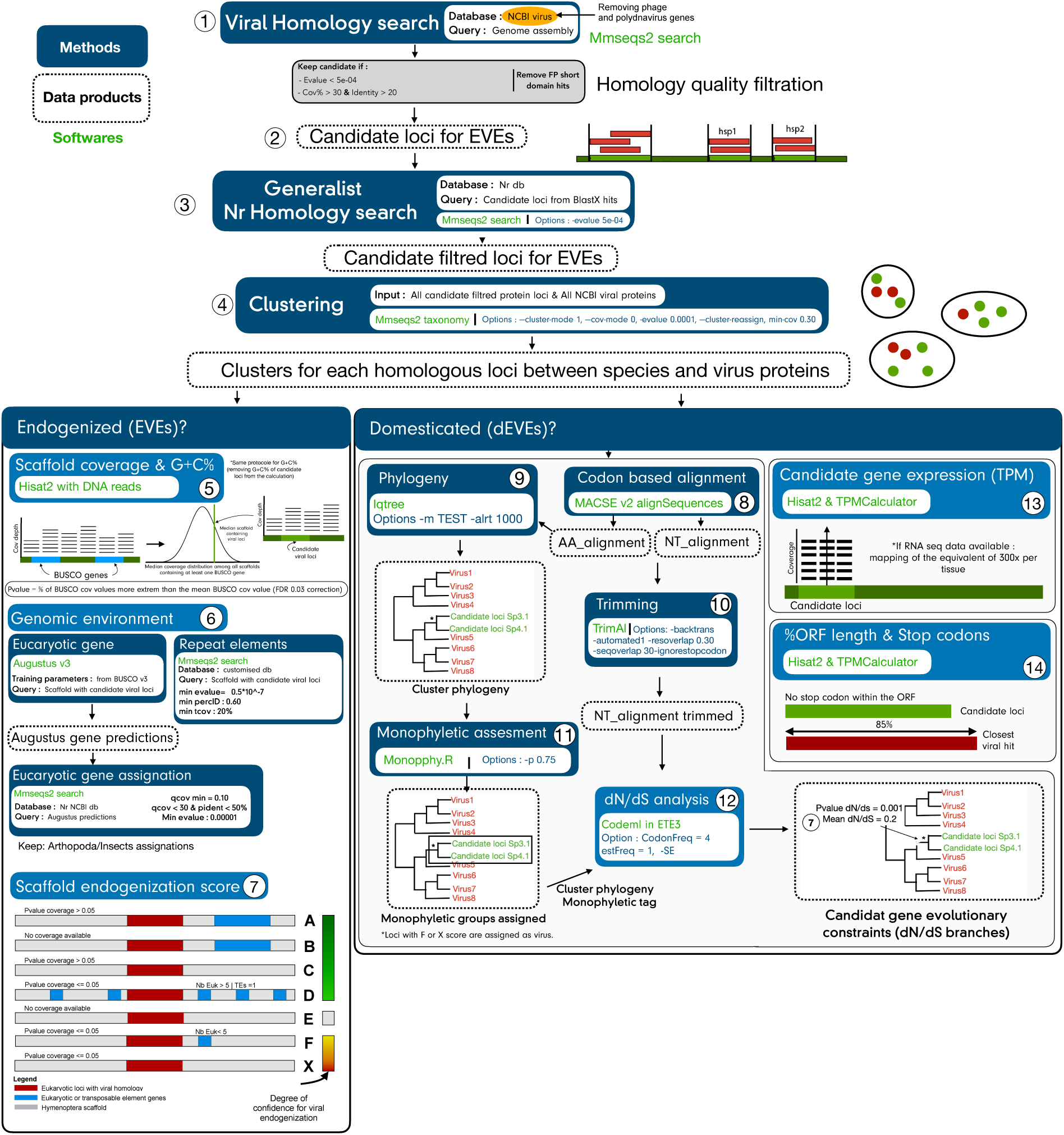
Simplified summary of the bioinformatics pipeline for the detection and validation of candidates for endogenization and domestication.

**Figure S14.**
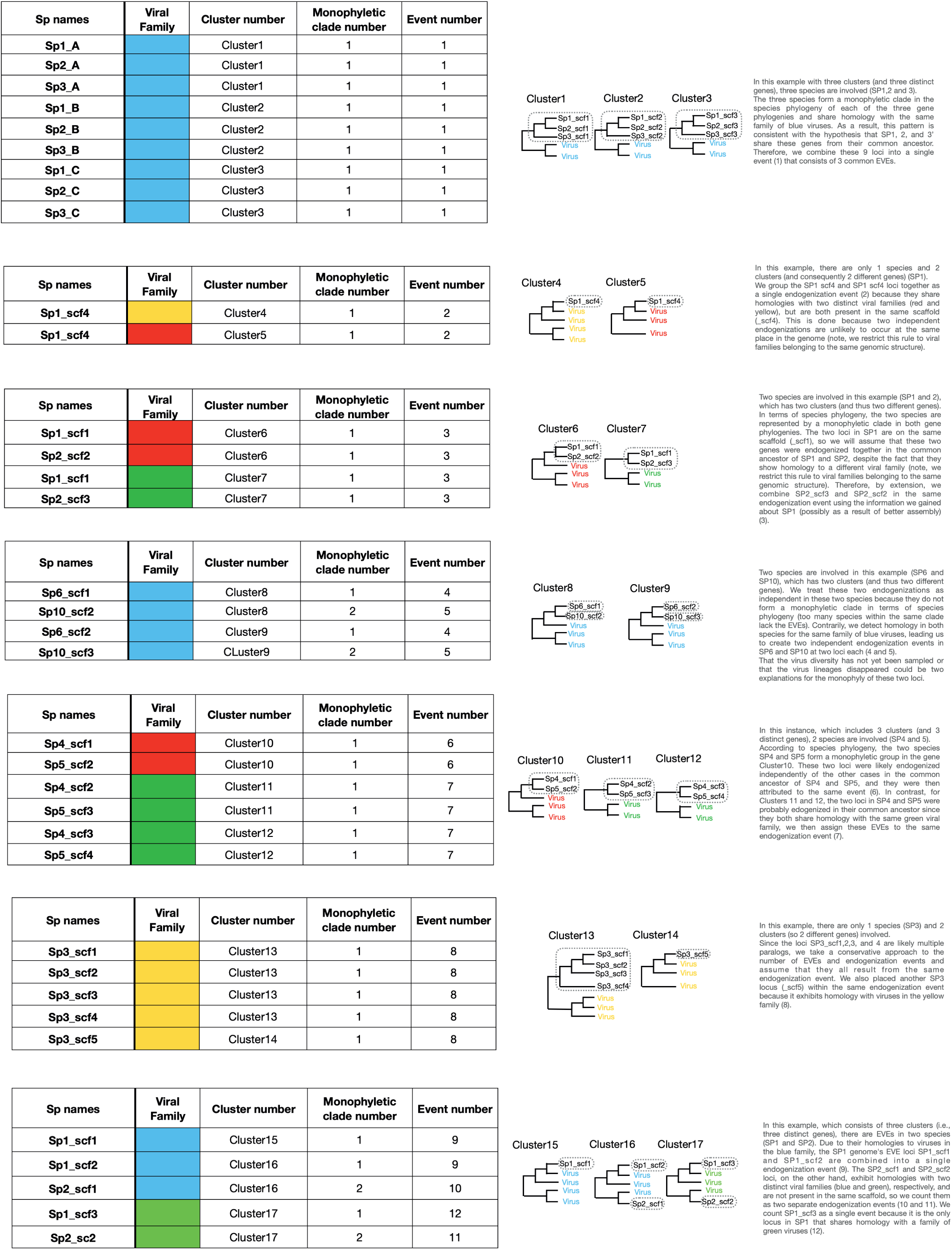
Canonical examples of endogenization events inferred by our pipeline. The column “Sp names” contains the species name, followed by the name of the scaffold in which the EVE has been identified. The “Viral family” column refers to the putative viral family that donated the EVEs. The column “Cluster number” corresponds to the number of the corresponding cluster phylogeny (thus the EVE phylogeny). The “Monophyletic clade number” column corresponds to the number of the monophyletic clade within a cluster (can be a single locus or multiple loci). The column “Event number” is the number given to single/multiple EVEs that derive from the same endogenization event.

**Table S1.** Summary statistics for control cases. The numerator indicates the numbers of EVEs or dEVEs inferred by our pipeline, and the denominator indicates the number of known EVEs for each case. Analysis on dN/dS was only possible when orthologs or paralogs were available. Controls Endogenous viral elements present in scaffolds probably belonging to the Hymenoptera genome are scored from A to D, and scaffolds probably belonging to free viruses are scored as F or X: (see details in Materials and methods). TPM (Transcripts per kilobase million) values were calculated via RNAseq read mapping when available in the databases (all RNAseq data sources can be found on the github repository under the name : RNA_seq_reads_mapped.txt). * In addition to the expected unique shared event concerning the *M. demolitor* and *C. vestalis* species, our pipeline inferred two additional events, each specific to one lineage. This was due to the fact that two genes were not detected by our pipeline as shared by *M. demolitor* and *C. vestalis*, either because they are effectively not shared (for 3 of them: HzNVorf118, like-*pif-4* (*19kda*), *fen-1*), or because of some false negative in one of the two lineage (for one of them:p33 (ac92)).

|  | *V. canescens* | *F. arisanus* | *C. vestalis* | *M. demolitor* | *L. boulardi* | *L. heterotoma* | *L. clavipes* | % Total |
| --- | --- | --- | --- | --- | --- | --- | --- | --- |
| ----- **NB EVEs**<sup>1</sup> | **36/40** | **42/47** | **20/21** | **18/25** | **12/13** | **12/13** | **12/13** | **88.4%** |
| Scaffold A | 36 | 42 | 18 | 16 | 11 | 6 | 8 | 90.13% |
| Scaffold B | 0 | 0 | 0 | 0 | 0 | 0 | 0 | 0% |
| Scaffold C | 0 | 0 | 0 | 0 | 1 | 6 | 4 | 7.24% |
| Scaffold- D | 0 | 0 | 2 | 2 | 0 | 0 | 0 | 2.63% |
| Scaffolds E-F-X | 0 | 0 | 0 | 0 | 0 | 0 | 0 | 0% |
| ----- **NB dEVEs** | **22/36** | **11/42** | **20/20** | **18/18** | **12/12** | **12/12** | **12/12** | **71.82%** |
| by dN/dS<sup>2</sup> | 3/3 | 0/1 | 16/20 | 15/18 | 12/12 | 12/12 | 12/12 | 70.39% |
| by TPM 1000 | 21 | 11 | 10 | 9 | 1 | 0 | 0 | 43.60% |
| ----- **Nb Events** | **1/1** | **1/1** | **3/1\*** |  | **1/1** |  |  | . |

1 When paralogs were detected on different scaffolds, the best scaffold score was used.
2 Analysis on dN/dS was only possible when orthologs or paralogs were detected (for example, in *V. canescens* this calculation was only possible for 3 genes having paralogs)

**Table S2.** Summary table. Clusters refers to the number of homologous clusters with one or more candidate Endogenous Viral Elements (EVEs). Raw EVEs is the raw number of EVEs (i.e. including all paralogs and orthologs) according to the scaffold categories from A to D. Nb EVEs with complete ORFs corresponds to the number of raw EVEs with an ORF starting with a methionine, without premature stop codons, and ending with a stop codon, distinguishing ORFs whose size is at least equal to or greater than 80% of that of the best viral hit. EVEs is the count of EVEs, i.e. counting the number of genes within each monophyletic group only once (as several EVEs may have undergone post-endogenization duplications or an EVE may be ancestrally acquired and shared by several species). Mean pident is the average of the percentage identity of all EVEs with the best viral hit, the number in brackets corresponds to the standard error. Nb EVEs pident and Nb EVE E-values 1e-20 correspond to the number of refined EVEs showing a hit with more than 80% identity and an E-value below 1e-20 with a viral protein, respectively. Domesticated EVEs (dEVEs) corresponds to the number of refined EVEs with either a dN/dS significantly less than 1 with a complete ORF and without a stop codon and/or a TPM value > 1000 with a complete ORF and without stop codon. Events corresponds to the number of endogenization events that may include one or more genes and involve one or more species. Shared Events is the number of endogenization events shared by at least two species. Viral families corresponds to the number of different putative viral families associated with the best viral hits. dEvents corresponds to the number of endogenization events presenting at least one dEVE.

|  | dsDNA | ssDNA | dsRNA | ssRNA | Total |
| --- | --- | --- | --- | --- | --- |
| Clusters | 166 | 9 | 18 | 36 | 229 |
| Raw EVEs<br>(A\|B\|C\|D) | 819<br>(534\|61\|102\|122) | 92<br>(40\|27\|6\|19) | 95<br>(45\|19\|26\|5) | 255<br>(129\|74\|34\|18) | 1261 |
| Compleat raw EVEs ORFs<br>(>80% best viral hit) | 694 (409) | 74 (39) | 76 (26) | 223 (76) | 1067 (550) |
| EVEs | 366 | 32 | 61 | 162 | 621 |
| Mean pident (sd) | 36.99 (15.35) | 36.91 (11.37) | 38.32 (13.27) | 33.30 (11.82) | 36.32 |
| EVE pidents >= 80 | 22 | 0 | 0 | 0 | 22 |
| EVE E-values 1e-20 | 430 | 29 | 67 | 126 | 652 |
| dEVEs | 112 | 9 | 12 | 38 | 171 |
| Events | 130 | 26 | 59 | 152 | 367 |
| shared Events | 17 | 1 | 5 | 13 | 36 |
| viral families/clades | 12 | 3 | 4 | 21 | 40 |
| dEvents | 47 | 10 | 12 | 38 | 107 |

**Table S3.** EVEs distribution according to putative non-segmented single-stranded RNA virus donor and their genomic position. For each EVE, we retrieved the position of the homologous ORF in the virus genome identified after a blastp search (first hit). The information on the position of the ORFs was retrieved either from [94], from the ICTV reports or manually after recovery of the viral assembly in NCBI, ORF annotation with getorf (ORF length min= 150bp) and blastp to confirm the position of the ORFs and their functions. The position of the ORF in the free-living virus genome is reported in the column “ORF position”, when the genome was incomplete, we inferred the position of the ORF with respect to the position of the homolog in the closest complete viral genome. The number of EVEs corresponds to the number of EVEs counting paralogs only once (i.e. counting only one EVE per species per gene cluster).

| Family | Closest viral blast hit | Protein ID | Protein Name | ORF position | Source | Nb EVEs |
| --- | --- | --- | --- | --- | --- | --- |
| *Artoviridae* | Pteromalus puparum peropuvirus | YP_009505431 | Glycosylated matrix protein M (U3) | 3 out 5 | ICTV | 19 |
| *Artoviridae* | Pteromalus puparum peropuvirus | YP_009505433 | RNA pol (L) | 5 out 5 | ICTV | 9 |
| *Artoviridae* | Beihai rhabdo like virus 1 | YP_009333442 | Phosphoprotein (U2) | 2 out 3 | ICTV | 1 |
| *Bornaviridae* | Estrildid finch bornavirus 1 | YP_009505428 | RNA pol (L) | 4 out 4 | Manually | 8 |
| *Chuviridae* | Hubei chuvirus like 1 | YP_009337906 | Nucleoprotein (N) | Incomplete (putative 3 out 3) | Manually | 24 |
| *Chuviridae* | Hubei coleoptera virus 3 | YP_009336865 | Nucleoprotein (N) | 2 out 3 | Shi et al 2016 | 6 |
| *Chuviridae* | Hubei chuvirus like 3 | YP_009337091 | Nucleoprotein (N) | 3 out 3 | Shi et al 2016 | 2 |
| *Chuviridae* | Hubei coleoptera virus 3 | YP_009336866 | RNA pol (L) | 3 out 3 | Shi et al 2016 | 2 |
| *Chuviridae* | Lonestar tick chuvirus | YP_009254003 | Nucleoprotein (N) | 3 out 3 | Manually | 1 |
| *Chuviridae* | Hubei odonate virus 11 | YP_009336948 | Nucleoprotein (N) | 3 out 3 | Shi et al 2016 | 1 |
| *Lispirividae* | Tacheng tick virus 6 | YP_009304417 | ORF1 | 1 out 5 | Manually | 5 |
| *Lispirividae* | Hubei rhabdo like virus 3 | YP_009336884 | Glyceraldehyde-3-phosphate dehydrogenase | 1 out 5 | Manually | 7 |
| *Lispirividae* | Hubei rhabdo like virus 3 | YP_009336887 | Glycoprotein (G) | 3 out 5 | Manually | 2 |
| *Lispirividae* | Hubei rhabdo like virus 3 | YP_009336889 | RNA pol (L) | 5 out 5 | Manually | 1 |
| *Nyamiviridae* | Midway nyavirus | YP_002905336 | Nucleoprotein (N) | 1 out 6 | ICTV | 8 |
| *Nyamiviridae* | Orinoco orinivirus | YP_009666283 | Nucleoprotein (N) | 1 out 4 | Manually | 8 |
| *Nyamiviridae* | Sierra Nevada nyavirus | YP_009044206 | Nucleoprotein (N) | 1 out 6 | ICTV | 5 |
| *Nyamiviridae* | Nyamanini nyavirus | YP_002905342 | Nucleoprotein (N) | 1 out 6 | ICTV | 4 |
| *Nyamiviridae* | Midway nyavirus | YP_002905331 | RNA pol (L) | 6 out 6 | ICTV | 1 |
| *Nyamiviridae* | Wenzhou Crab Virus 1 | YP_009304556 | Nucleoprotein (N) | 1 out 4 | Manually | 1 |
| *Nyamiviridae* | Beihai rhabdo like virus 3 | YP_009666292 | RNA pol (L) | 5 out 5 | Shi et al 2016 | 1 |
| *Rhabdoviridae* | Wugan Ant virus | YP_009304559 | RNA pol (L) | Incomplete (putative 5 out 5) | Manually | 2 |
| *Rhabdoviridae* | Wuhan insect virus 7 | YP_009301743 | RNA pol (L) | 5 out 5 | Manually | 1 |
| *Rhabdoviridae* | Sanxia Water Strider Virus 5 | YP_009289351 | Glycoprotein (G) | 4 out 5 | Manually | 1 |
| *Rhabdoviridae* | Muscina stabulans sigmavirus | YP_009664711 | RNA pol (L) | Incomplete (putative 5 out 5) | Manually | 1 |

## Notes

https://github.com/BenjaminGuinet/Supplementary-Endoparasitoidism_promotes_viral_domestication

https://github.com/BenjaminGuinet/Viral_domestication_finder_V2

### Competing Interest Statement

The authors have declared no competing interest.

### Summary of Updates

Update of the Hymenoptera phylogenetic analysis datation to correct a minor error in the choice of amino acid sites.

